# Evolutionary time best explains the latitudinal diversity gradient of living freshwater fish diversity

**DOI:** 10.1101/668079

**Authors:** Elizabeth Christina Miller, Cristian Román-Palacios

**Affiliations:** Department of Ecology and Evolutionary Biology, University of Arizona, Tucson, Arizona 85721, U.S.A.; School of Aquatic and Fishery Sciences, University of Washington, Seattle, Washington 98105, U.S.A.

**Author notes:** Authors contributed equally to this work. **Elizabeth Miller** is an NSF Postdoctoral Fellow at the University of Washington. She received her Ph.D. from the University of Arizona. She is an ichthyologist interested in explaining how speciation, extinction, and colonization vary to produce modern biodiversity patterns. **Cristian Román-Palacios** is a doctoral candidate at the Department of Ecology and Evolutionary Biology at the University of Arizona. He studies large-scale patterns of biodiversity from an Eco/Evo perspective and examines the effects of climate change on species survival.

**Keywords:** diversification rates, freshwater fishes, Generalized Additive Models, latitudinal diversity gradient, species richness, time-for-speciation

## Abstract

**Aim:** The evolutionary causes of the latitudinal diversity gradient are debated. Hypotheses have ultimately invoked either faster rates of diversification in the tropics, or more time for diversification due to the tropical origins of higher taxa. Here we perform the first test of the diversification rate and time hypotheses in freshwater ray-finned fishes, a group comprising nearly a quarter of all living vertebrates.

**Location:** Global.

**Time period:** 368–0 mya.

**Major taxa studied:** Extant freshwater ray-finned fishes.

**Methods:** Using a mega-phylogeny of actinopterygian fishes and a global database of occurrence records, we estimated net diversification rates, the number of colonizations and regional colonization times of co-occurring species in freshwater drainage basins. We used Generalized Additive Models to test whether these factors were related to latitude. We then compared the influence of diversification rates, colonization numbers, colonization times and surface area on species richness, and how these factors are related to each other.

**Results:** While both diversification rates and time were related to richness, time had greater explanatory power and was more strongly related to latitude than diversification rates. Other factors (basin surface area, number of colonizations) also helped explain richness but were unrelated to latitude. The world’s most diverse freshwater basins (Amazon, Congo rivers) were dominated by lineages with Mesozoic origins. The temperate groups dominant today arrived near the K-Pg boundary, leaving comparatively less time to build richness. Diversification rates and colonization times were inversely related: recently colonized basins had the fastest rates, while ancient species-rich faunas had slower rates.

**Main conclusions:** We concluded that time is the lead driver of latitudinal richness disparities in freshwater fish faunas. We suggest that the most likely path to building very high species richness is through diversification over long periods of time, rather than diversifying quickly.

## 1 INTRODUCTION

Species richness decreases from the equator to the poles. The latitudinal biodiversity gradient has been called the Earth’s first-order biodiversity pattern due to its pervasiveness across groups and geologic time (Hillebrand, 2004). There are only three processes that can directly change regional species richness: in-situ speciation, local extinction, and dispersal (Ricklefs, 1987; Roy & Goldberg, 2007). While numerous ecological and evolutionary hypotheses have been proposed to explain the latitudinal biodiversity gradient (Mittelbach et al., 2007), it is clarifying to first determine how these three core processes change with latitude. Other factors such as area or productivity may also influence richness (Tedesco et al., 2012), but these factors must change richness indirectly by acting on speciation, extinction, and/or dispersal.

There has been great interest in comparing diversification rates across phylogenies in recent years, due to the confluence of the construction of large time-calibrated molecular phylogenies (e.g. Jetz, Thomas, Joy, Hartmann, & Mooers, 2012; Rabosky et al., 2018) and the increasing complexity of models of diversification (e.g. Rabosky, 2014). A growing number of analyses across groups are revealing that speciation and/or net diversification rates are similar among latitudes (general review in Schluter & Pennell, 2017; Jansson, Rodríguez-Castañeda, & Harding, 2013; ants: Economo, Narula, Friedman, Weiser, & Guénard, 2018; birds: Weir & Schluter, 2007; Jetz et al., 2012; Rabosky, Title, & Huang, 2015), or even faster in high latitudes (angiosperms: Igea & Tanentzap, 2019; deep-sea invertebrates: O’Hara, Hugall, Woolley, Bribiesca-Contreras, & Bax, 2019; mammals: Morales-Barbero, Gouveia, & Martinez, 2020; marine fishes: Rabosky et al., 2018; Miller, Hayashi, Song, & Wiens, 2018). These studies raise the question: if diversification rates do not explain spatial differences in richness then what does?

A potential resolution to this question is to compare the relative importance of colonization history and diversification rates for explaining species richness (Stephens & Wiens, 2003). This distinction is important because while we may identify in-situ speciation as the dominant process generating biodiversity in a region (Tedesco et al., 2012), similar richness disparities may be produced either through faster speciation or speciation over longer periods of time (Figure 1; Wiens, 2012). For example, tropical marine fishes have modest speciation rates (Rabosky et al., 2018), but the tropics still have high richness due to the combination of early and frequent colonization (Miller et al., 2018). Similarly, time matters for explanations involving asymmetrical dispersal rates. Frequent colonization may not lead to high richness if this trend is recent, such that new colonists have not had much time to diversify (Miller et al., 2018). The time-for-speciation effect may also imply spatial variation in extinction pressure. One reason that a region may be dominated by young lineages is that older lineages went extinct (Miller & Wiens, 2017). The influence of time on species richness is an idea with deep historical roots (Wallace, 1878; Willis, 1922; Wiens & Donoghue, 2004; Fine & Ree, 2006; Jablonski, Roy, & Valentine, 2006; Mittelbach et al., 2007). While researcher attention on time as an explanation for the latitudinal diversity gradient has waxed and waned over the years (Stephens & Wiens, 2003), recent studies that have simultaneously inferred colonization and diversification history have found a strong role for time in explaining richness patterns (Jansson et al., 2013; Miller et al., 2018; Economo et al., 2018; O’Hara et al., 2019; Li & Wiens, 2019).

**FIGURE 1.**
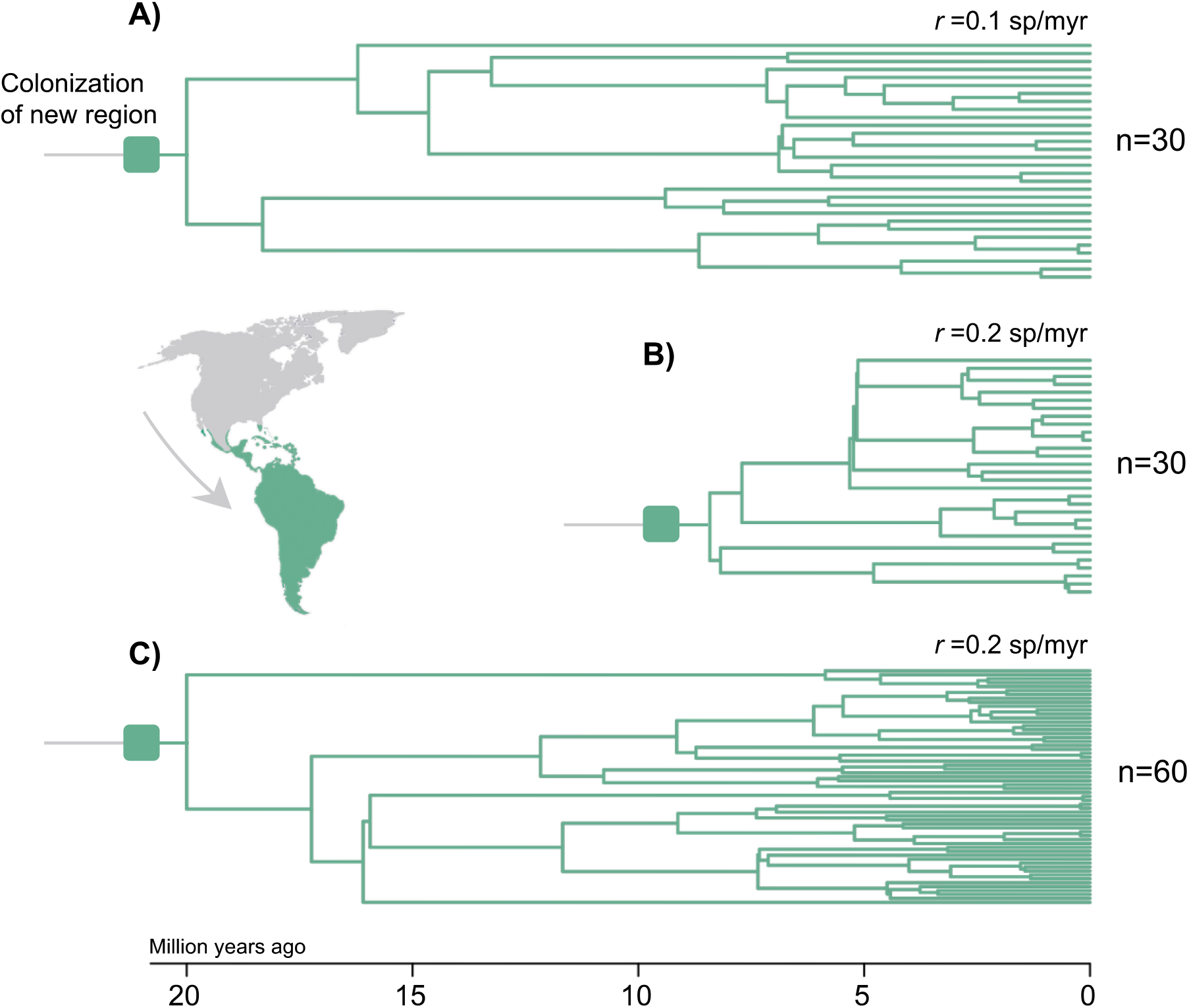
Schematic illustrating alternative pathways to building species richness through in-situ speciation. In all three examples, a lineage colonizes a region and diversifies with no further immigration or emigration. In (**A**), the lineage colonizes the region relatively early and diversifies at a modest rate. In (**B**), the lineage colonizes later, but diversifies at a faster rate and achieves the same species richness as (**A**). In (**C**), the lineage colonizes relatively early and also diversifies quickly, achieving high richness. While diversification rates are constant in these simple examples, these scenarios are also applicable to cases where rates change through time (Pontarp & Wiens, 2017).

A related hypothesis, that regions with more species have greater ecological limits, ultimately implies that speciation, extinction or dispersal rates change in association with a carrying capacity of species (Rabosky, 2009; Hurlbert & Stegen, 2014). Thus, a carrying capacity influences richness indirectly by acting on speciation, extinction or dispersal, just as area or climate influences richness. Ecological limits are not inconsistent with either the rate or time hypotheses. For example, regions with low carrying capacities can take longer to be colonized in simulations, showing that ecological limits can modulate richness through the time-for-speciation effect (Pontarp & Wiens, 2017). If a region was colonized early in the history of a clade, it is generally expected to contain more species than recently colonized regions due to greater time allowed for speciation, even if speciation rates have slowed through time (Pontarp & Wiens, 2017). Species richness will increase over time as long as speciation rates are non-zero and exceed extinction rates (Machac, 2020).

Freshwater fishes represent nearly a quarter of all vertebrate species (Cavin, 2017) and are major components of ecosystems in both tropical and temperate latitudes (Berra, 2001). A body of work has demonstrated correlations with freshwater fish species richness and factors including productivity and area (Oberdorff, Guégan, & Hugueny, 1995; Smith, Badgley, Eiting, & Larson, 2010), in-situ speciation (Tedesco et al., 2012) and recent historical events such as Quaternary sea-level changes (Oberdorff, Guégan, & Hugueny, 1997; Tedesco, Oberdorff, Lasso, Zapata, & Hugueny, 2005; Leprieur et al., 2011; Dias et al., 2014). At least three questions remain about what drives species richness in freshwater habitats. First, while in-situ speciation (cladogenesis) is clearly important as inferred from endemism in species-rich basins (Tedesco et al. 2012), is high species richness caused by faster rates of speciation or more time for speciation due to earlier colonization (Figure 1)? Second, since area cannot directly change species richness, in what potential ways is the species-area relationship related to latitudinal trends in speciation and colonization? Third, recent events seem to impact local richness within a continent, but differences in richness among continents remain (Oberdorff et al., 1997; Tedesco et al., 2005; Dias et al., 2014). How did these continental differences in richness form? We attempt to answer these questions herein.

Here we test whether variation in diversification rates or time-for-speciation best explains global diversity patterns in freshwater fishes, especially the latitudinal diversity gradient. Freshwater fishes have features that are conducive to testing both diversification rates and time hypotheses. First, living freshwater fishes represent a wide range of ages, with some radiations diversifying during the Mesozoic or earlier (Briggs, 2005; Capobianco & Friedman, 2018) and others only during the most recent glaciation cycles (Seehausen & Wagner, 2014). Second, freshwater fishes have low dispersal ability, and therefore their systematics likely retain signatures of regional events (Lavoué, 2016; Cavin, 2017; Capobianco & Friedman, 2018). Our study capitalizes on the aggregation of natural history observations and genetic data over many years (Tedesco et al., 2017; Rabosky et al., 2018), allowing us to make comparisons at a broad spatial and temporal scale under a common phylogenetic framework.

## 2 METHODS

Additional details are given in the Supporting Information.

### 2.1 Occurrence and phylogenetic data

Expert-vetted occurrence records of freshwater actinopterygian species were assembled by Tedesco et al. (2017). Occurrence records were available for 3,119 drainage basins among six biogeographic regions. This dataset also reported surface area (Figure S1), median latitude and longitude of each basin. We removed non-native and uncertain records. Altogether, occurrence records from 14,947 species of freshwater fishes were used to estimate species richness of basins. Drainage basins with species records covered 80% of the Earth’s land surface overall and ranged from 70% of the surface of the Indo-Malay region to 90% of the surface of the Afrotropics (Tedesco et al., 2017). Species coverage for major freshwater fish groups ranged from 61% of Anabantiformes to 93% of Characiformes.

For biogeographic and diversification-rate analyses, we used the maximum likelihood time-calibrated molecular phylogeny of actinopterygians constructed by Rabosky et al. (2018; see also http://fishtreeoflife.org). This phylogeny includes 11,638 species with genetic data (36.9% of known ray-finned fishes).

### 2.2 Obtaining diversification rates and colonization times for basins

To estimate diversification rates for each drainage basin, we used three types of branch-associated measures. We used tip-based net diversification rates calculated using BAMM v.2.5.0 (Rabosky, 2014): three independent runs under a constant-rate model of diversification, and three runs under a time-varying model. We also used tip-based estimates of the DR statistic (Jetz et al., 2012). Unlike BAMM, the DR statistic was calculated from phylogenies with all unsampled species grafted using taxonomic constraints (n=31,516 species). We note that DR tip rates better approximate speciation rates than net diversification rates in comparison to rates from BAMM (Title & Rabosky, 2019). For each measure of tip-based rates, we took the mean of rates among co-occurring species in each basin.

The “time-for-speciation” effect represents the time allowed for in-situ diversification since a lineage colonized a region (Figure 1; Stephens & Wiens, 2003). Note that our preferred terminology is to use “dispersal” to refer to the general and bi-directional process of movement among regions, “colonization” as the addition of new lineages to a focal region as a result of dispersal, and “time-for-speciation” as the time elapsed between colonization and the present (Stephens & Wiens, 2003; Hua & Bromham, 2020). To measure time-for-speciation, we must estimate the amount of time a lineage has been present in the location of interest (Figure 1, Figure S2). One major challenge to estimating colonization times at the local scale is that the computation time of biogeographic models scales exponentially with the number of possible ranges (Matzke, 2014). Modelling dispersal among >3,000 drainage basins is unfeasible using phylogenetic approaches at present. To overcome this challenge, we instead modelled dispersal among continental regions. We then used the mean and median regional colonization time associated with species present in each drainage basin. This approach to measuring spatial variation in time-for-speciation is analogous to the grid-cell approach often used to detect spatial variation in diversification rates (Jetz et al., 2012; Rabosky et al., 2018; Machac, 2020). By focusing on regions, our colonization time estimates should also be more robust to past range shifts since regions change less over time than individual rivers and lakes (Hoorn et al., 2010).

We first fit a dispersal-extinction-cladogenesis model (DEC; Ree & Smith, 2008) using the R package ‘BioGeoBEARS’ v.1.1 (Matzke, 2014; additional details in Extended Methods, Supporting Information). We used the maximum likelihood phylogeny including species with genetic data only (Rabosky et al., 2018), because semi-random grafting of unsampled species is inappropriate for comparative methods that model the evolution of traits associated with the tips (Rabosky, 2015). We removed 139 tips that were unsuitable for biogeographic reconstructions, leaving 11,499 species. Our analysis included six continental regions following Tedesco et al. (2017): the Neotropics, Afrotropics, Indo-Malay, Australasia, Nearctic, and Palearctic. Species restricted to marine environments were coded as occurring in a seventh “marine” region. Though not our focus, these species are needed to inform the timing of colonization of freshwater regions from the marine realm (Betancur-R, Ortí, Stein, Marceniuk, & Pyron 2012; Betancur-R, Ortí, & Pyron, 2015; Rabosky, 2020). Our model was time-stratified to apply constraints on dispersal in accordance with changing connectivity of continents. We applied the following constraints over six time bins spanning the root (∼368 mya) to the present (following Toussaint, Bloom, & Short, 2017): dispersal between adjacent regions was not constrained (i.e. the probability of dispersal between adjacent regions was scaled by 1); dispersal between regions separated by a small marine barrier was scaled by 0.75; dispersal between regions separated by another landmass was scaled by 0.50; and dispersal between regions separated by a large marine barrier was scaled by 0.25 (more details in Table S1).

After model fitting, in order to identify individual colonization events and visualize uncertainty in these dates, we simulated 100 biogeographic stochastic maps (Dupin et al., 2016). Each individual simulation is a realized history that is possible given the model and data, including the time and location on the branches for biogeographic events. Averaging over all of these simulations will approximate the ancestral state probabilities calculated by the model. We used these simulations to estimate the time-for-speciation associated with each drainage basin. To do this, we traced each individual species back in time to the location on the branch reconstructed as the colonization of the region(s) it inhabits. Note that it is possible for this event to precede the crown age of recognizable clades (such as orders), especially at this large phylogenetic scale, since dispersal can happen at any time along a phylogeny (Hua & Bromham, 2020). See Figure S2 for an illustration for how these times were obtained. For each species we took the mean time of this event across the 100 stochastic maps. The amount of time-for-speciation associated with each basin was estimated as the mean and median colonization time among co-occurring species.

The number of colonizing lineages can also predict richness (Miller et al., 2018). Estimating the number of colonizations to individual drainage basins has the same challenge as estimating time associated with basins (model constraints; see above). Instead, we counted the number of independent colonizations to the major region represented among co-occurring species in each basin. We used the mean of this count among 100 stochastic maps in analyses.

### 2.3 Comparing predictors of local richness

To sum, we first tested how basin richness, diversification rates, time-for-speciation, and surface area change with latitude and longitude. Second, we tested whether diversification rates and time-for-speciation each separately predict local richness. Third, we compared the relative support for diversification rates and time-for-speciation for predicting richness, with and without area as a covariate. Fourth, we tested how diversification rates and time-for-speciation are related to each other. Fifth, we tested whether variation in area with latitude could explain our results. Sixth, we tested whether the number of colonizations was related to richness, latitude, time or diversification rates.

We fit Generalized Additive Models (GAMs) to examine the change in species richness, net diversification rates, and time-for-speciation with latitude, longitude, and both (i.e. the interaction between longitude and latitude). We considered longitude as well as latitude because species richness also varies strongly within the tropics (Figure 3A). We fit univariate GAMs between predictor (latitude or longitude) and response variables (species richness, net diversification rates, or time-for-speciation). GAMs were made spatially explicit by including the interaction between longitude and latitude of each basin in the smooth term (i.e. s(long, lat)). This approach for explicitly accounting for geography in GAMs was first proposed by Brumback & Rice (1998), with further details presented in Kammann & Wand (2003), Hefley et al. (2016), and Wood (2017). Univariate GAMs were fit using the gam function in the ‘mgcv’ package in R (base; R Core Team, 2008; Wood, 2011). To assess the direction of the relationships we performed Spearman’s rank correlation tests between latitude and species richness, net diversification rates, time-for-speciation and surface area. We also fit a GAM to confirm whether surface area was related to species richness.

Next, we fit spatially explicit GAMs to understand the relative importance of diversification rates and time-for-speciation for explaining local richness. Again, to account for spatial autocorrelation, we included a smoother term that summarized the interaction between latitude and longitude in each basin (e.g. s(long, lat) in GAMs). We first analyzed the relationships between basin richness and each predictor alone. Then, we included both variables as predictors of richness. We compared the fit of four models using AIC values: a null model that assumed species richness to be constant (“null model”); a model where species richness depended on diversification rates (“div model”); a model where species richness depended with time-for-speciation (“time model”); and a model where species richness depended on both diversification rates and time-for-speciation (full model). We then estimated the amount of deviance in richness explained by each predictor (details in the Extended Methods, Supplementary Information). We fit this set of four models for each combination of diversification rate estimate (seven alternatives) and estimate of basin-level colonization times (two alternatives). In addition, we repeated these analyses while also controlling for the effect of surface area (i.e. species-area scaling). For each combination of diversification rate and colonization times, we fit the same four models described above with the addition of surface area as a covariate.

We examined how two predictors of richness (diversification rates, time-for-speciation) interact with each other. For example, if diversification rates and time-for-speciation are jointly responsible for producing the most-diverse faunas, then one would expect these basins to harbor rapidly-diversifying lineages that colonized a long time ago (Figure 1). Alternatively, some basins may have fast diversification rates while others were colonized early. We fit spatially explicit GAMs with time-for-speciation as the predictor of diversification rates. We also performed Spearman’s rank correlation tests between diversification rates and time-for-speciation to quantify the strength and direction of the association between these variables.

Following this, we fit spatially explicit GAMs to assess the possibility that a covariation between surface area and latitude was responsible for trends in time-for-speciation and diversification rates. We fit a set of three GAMs for each target variable (time-for-speciation and diversification rates). First, we fit a full model including the additive effects of area and latitude in explaining spatial patterns in either time-for-speciation or diversification rates. Next, we fit two additional models with changes in the predictor being explained by either latitude or area alone. To compare the relative importance of latitude and surface area for explaining diversification rates and time-for-speciation, we estimated the change in deviance from the exclusion of each predictor compared to the full model.

Finally, we fit spatially explicit GAMs to test if the number of independent colonizations was related to richness, and how this number is related to latitude, diversification rates and time. We also fit linear regressions and used the slope to describe the nature of the relationship between these variables. For example, it is possible that the number of colonizations influences richness but only in some groups of basins (e.g. young or slowly-diversifying basins).

### 2.4 Assessing sensitivity of colonization time estimates

Several freshwater clades are known to have extinct marine members (Betancur-R et al., 2015). Colonization times associated with these groups may be overestimated without accounting for marine ancestry erased by extinction. To assess this, we also performed ancestral range reconstructions using an alternative time-calibrated phylogeny containing 1,582 living and 240 extinct ray-finned fishes (Betancur-R et al., 2015). We used the literature to assign freshwater fossil species to continental regions (Table S2). Biogeographic analyses were performed as above (see also Supporting Information). We compared the mean and range of colonization times inferred from biogeographic stochastic mapping for major freshwater fish clades between the two phylogenies. From this comparison, we found that colonization times associated with early-diverging (non-teleost) clades were overestimated using the Rabosky et al. (2018) phylogeny. We removed the four living non-teleost orders (Polypteriformes, Acipenseriformes, Lepisosteiformes and Amiiformes) from our basin-level dataset to quantify their impact on basin-specific diversification-rate and time-for-speciation estimates.

## 3 RESULTS

Richness of freshwater actinopterygian fishes among basins reflects the latitudinal diversity gradient found in other major groups (Figure 2A; Spearman’s rank correlation between local richness and latitude: rho=-0.27, P<0.001). However, latitude alone only explains ∼9% of the variance in species richness across basins (GAM, R^2^=0.092, P<0.001; Figure 2A; Table S3). Longitude alone had similar explanatory power (Figure 2B; GAM, R^2^=0.10, P<0.001). The interaction between longitude and latitude explains a higher proportion of the variance in species richness (R^2^=0.26, P<0.001). These results appear to reflect the strikingly high richness in the Neotropics relative to other tropical regions. For example, the Amazon basin contains about twice as many known species as the world’s second-most rich basin, the Congo (2,968 vs 1,554 species, respectively; Figure 3A). In sharp contrast, the most species-rich basin in the Australasian tropics is the Ramu River of New Guinea with 179 species. Therefore, while we found a latitudinal gradient in freshwater fish richness, much variation in species richness was unrelated to latitude.

**FIGURE 2.**
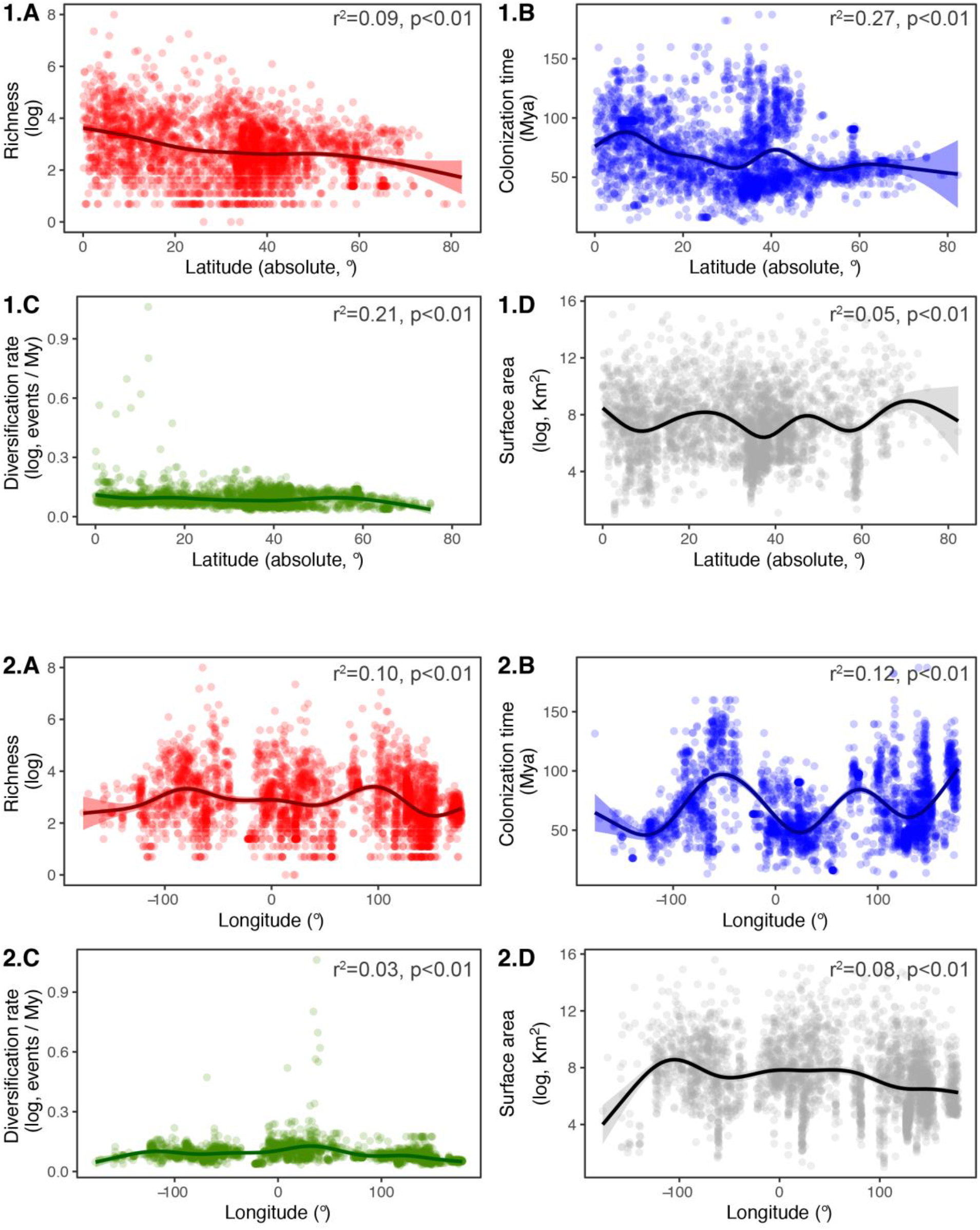
Latitudinal (**1**) and longitudinal (**2**) gradients of species richness (**A**), mean colonization times (**B**), mean net diversification rates (**C**), and surface area (**D**) of freshwater drainage basins. Species richness and surface area is derived from Tedesco et al. (2017). Diversification rates shown here were estimated using BAMM under a time-constant rates model; values represent the mean tip-associated values of species found in each basin (Rabosky et al., 2018). Colonization timing of biogeographic regions was inferred from ancestral range reconstructions (Matzke, 2014); values represent the mean regional colonization time for species in each basin. For full GAM results see Table S3.

**FIGURE 3.**
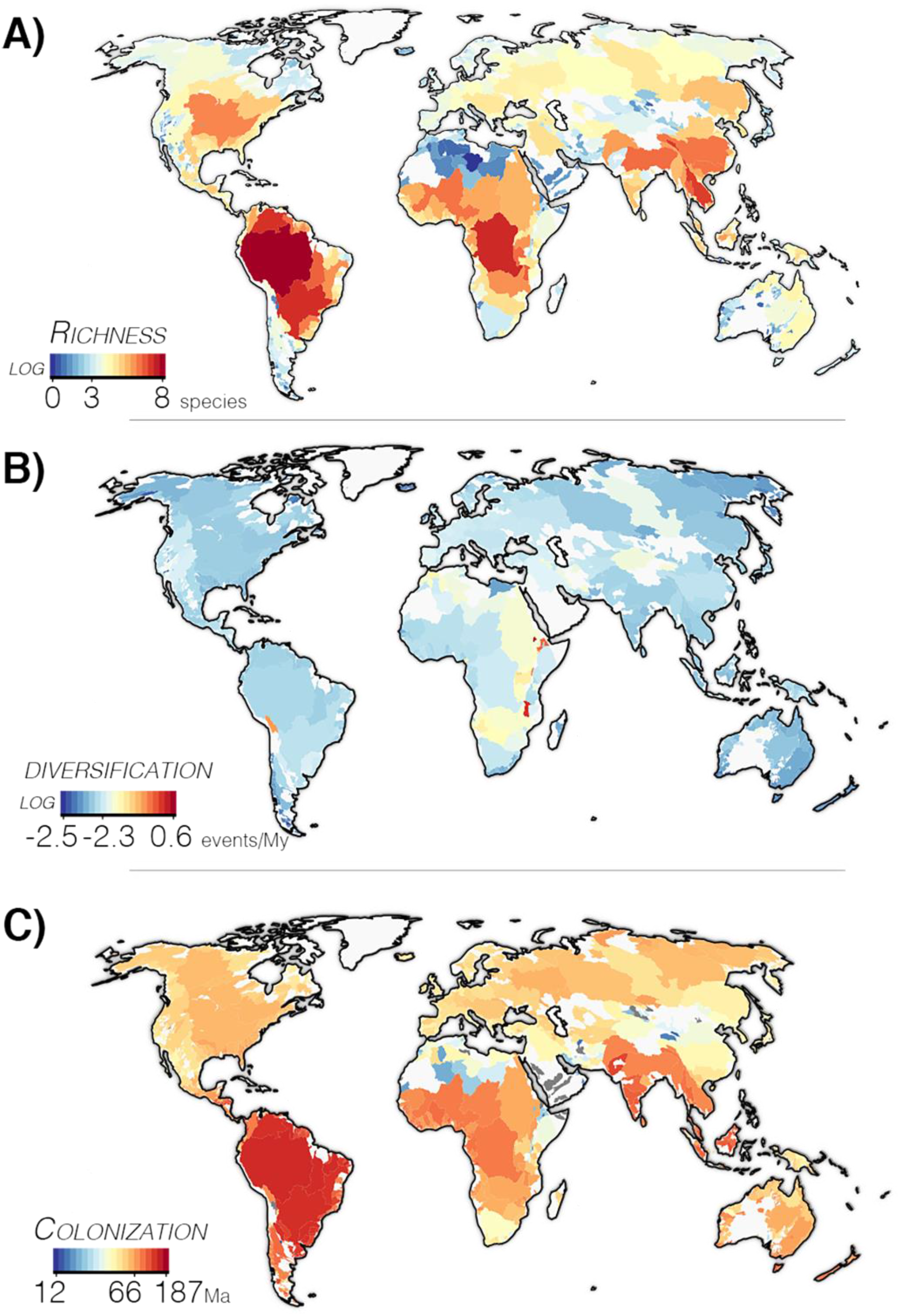
Results for spatially explicit GAMs for predictors of species richness of freshwater drainage basins. Geographical distribution of global freshwater fish (**A**) richness, (**B**) net diversification rates, and (**C**) time-for-speciation. Species richness of basins is based on occurrences from Tedesco et al. (2017). BAMM-estimated rates of net diversification calculated under a time-constant rates model (Rabosky et al., 2018) are shown here. Colonization times of biogeographic regions were inferred from ancestral range reconstructions (Matzke, 2014; see Extended Methods). Values of net diversification rates and colonization times represent the means among co-occurring species in each basin. Species richness and diversification rates are log-transformed. For full GAM results see Tables S4, S5; for bivariate relationships see Figure S3.

Next, we tested for latitudinal trends in diversification rates, time-for-speciation, and surface area. Latitude was significantly related to diversification rates (rho=-0.11–0.33, R^2^=0.205–0.212, P<0.001; Figures 2A, 3B; Table S3). This trend may have been influenced by outlier basins with very high rates (e.g. Lake Malawi and Lake Titicaca; Figure 3B). We also found a latitudinal trend in time-for-speciation, in which ancient colonizations were typical of low-latitude basins (median colonization time: R^2^*=*0.253, rho=-0.289, P<0.001; mean colonization time: R^2^*=*0.271, rho=-0.232, P<0.001; Figures 2A, 3C; Table S3). As expected, species richness was positively related to surface area of drainage basins (R^2^=0.242, P<0.001; Figure S1A). However, surface area was poorly related to latitude (R^2^=0.054, P<0.001; Table S3; Figure 2A; Figure S1B). Therefore, the latitudinal diversity gradient is potentially explained by geographic trends in time-for-speciation and diversification rates, but is unlikely to be explained by a relationship between present-day area and latitude.

We examined the individual effects of diversification rates and time-for-speciation for explaining richness patterns in general (not just in association with latitude). Both diversification rates and time-for-speciation were significantly related to richness and explained a similar portion of its variance globally (diversification rates: R^2^=0.338–0.364, all P<0.001; time-for-speciation: R^2^=0.317–0.326, all P<0.001; Table S4; Figure S3). Adding surface area as a covariate increased the variation in richness explained by GAMs (diversification rates: R^2^=0.534–0.554, all P<0.001; time-for-speciation: R^2^=0.527–0.534, all P<0.001; Table S5), reflecting the positive relationship between area and richness (Figure S1).

Our finding that both diversification rates and time-for-speciation influence species richness does not necessarily mean that these two processes have synergic effects in each basin (Figure 1). We used GAMs to test for a relationship between basin-level diversification rates and colonization times. We found that colonization times explain 39–42% of variance in diversification rates (all P<0.001; Table S6). Importantly, we found a negative relationship between diversification rates and colonization times, such that diversification rates tend to be higher in recently-colonized basins (Figure 4; rho=-0.13–-0.10, all P<0.001; Table S6). These results suggest that basins with fast diversification rates are often distinct from those with early colonizations (Figure 1).

**FIGURE 4.**
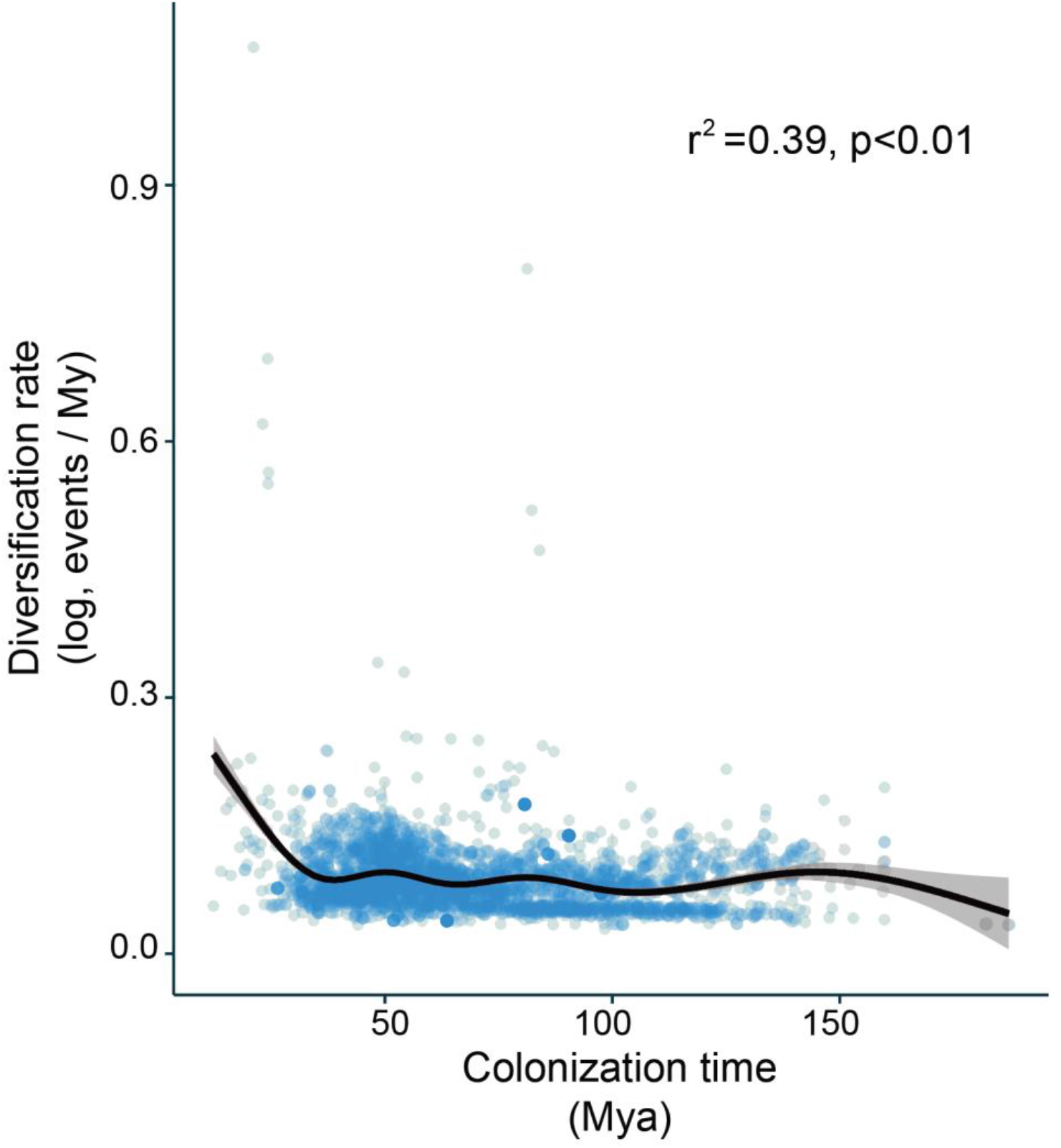
The negative relationship between net diversification rates and colonization times (GAM R^2^=0.393; Spearman’s rho=-0.125, both P<0.001). Values represent the means among co-occurring species in each drainage basin. BAMM-estimated rates of net diversification calculated under a time-constant rates model (Rabosky et al., 2018) are shown here. Results under a time-varying rates model were similar (Table S6). Colonization times of biogeographic regions were inferred from ancestral range reconstructions (Matzke, 2014; see Extended Methods).

We then compared the relative importance of diversification rates and time-for-speciation for explaining species richness. We found that time-for-speciation contributed 2.3–6.1 times more to species richness patterns than diversification rates (based on deviance values for alternative GAMs; Table S4). Time-for-speciation contributed 1.4–4.6 times more than diversification rates based on models that included surface area (Table S5).

We then asked whether latitudinal trends in time-for-speciation and diversification rates could be responding to a covariance between latitude and area. We found that latitude contributes 15 times more to variation in diversification rates than does surface area (Table S7). Latitude contributes 119 times more to variation in time-for-speciation than surface area. This suggests that latitudinal trends in diversification rates and time-for-speciation (Tables S4, S5) are unlikely to be explained by latitudinal trends in surface area of drainage basins.

Finally, we found a strong positive relationship between basin richness and the number of independent colonization events represented among the basin’s fauna (linear regression: R^2^=0.622, P<0.001, slope=1.13; Table S8; Figure S4). However, the number of colonizations was unrelated to latitude or diversification rates (latitude: R^2^<0.01, P=0.11; diversification: R^2^<0.01, P=0.15). While the number of colonizations is important for explaining richness in general, this number cannot explain latitudinal trends in richness. We also found that the number of colonizations was weakly but inversely related to the mean time of colonization (R^2^=0.067, P<0.001, slope=-9.995). This suggests that, like diversification rates, the number of colonizations may be most relevant for explaining richness among recently colonized basins.

To sum, diversification rates, time-for-speciation, surface area, and the number of colonizations were each significantly related to species richness. However, time-for-speciation was more strongly related to latitude than other variables. These results are reflected by the spatial patterns in richness and colonization times presented in Figure 3. While species-rich tropical basins such as the Amazon and Congo had a mean colonization time during the Mesozoic, 95.4% of basins in the Nearctic and Palearctic had mean colonization times after the K-Pg boundary (66 mya; Figure 3C; Table A4). Colonization times and diversification rates were inversely related. This suggests that the most species-rich basins are so because of more time for diversification, not faster diversification. Differences in diversification rates and the number of colonizations were also important for explaining richness patterns but were more relevant among recently colonized (and relatively depauperate) basins. Note that these results were robust to alternative estimates of diversification rates (BAMM time-constant, BAMM time-varying, and DR) and whether we used the mean or median colonization time among co-occurring species (Tables S3–S6).

We compared our colonization time estimates to those using an alternative phylogeny that also included fossils (Betancur-R et al., 2015). Aside from non-teleosts, colonization patterns for major freshwater fish lineages were generally congruent between the two phylogenies (Rabosky et al., 2018; Betancur-R et al., 2015). Despite uncertainty in colonization times in the ancestral reconstructions, the order of events was consistent such that colonization of tropical regions generally preceded colonization of extratropical regions (Extended Results 1; Figure S5). For more details see Extended Results 1.

The addition of fossils, especially marine members of families now restricted to freshwater, suggested that colonization times estimated for early-diverging fishes were overestimated (Extended Results 1; Figure S5). However, removing non-teleosts had a negligible effect on diversification rates globally (mean of 0.24% faster; Extended Results 2). Removing these species resulted in slightly younger mean colonizations globally (mean of 1.80% younger). Most basins affected (76%) were in the Nearctic and Palearctic. Results based on teleosts alone would widen the difference between tropical and extratropical time-for-speciation, in line with our conclusions based on all actinopterygians.

## 4 DISCUSSION

The three processes that directly change species richness are in-situ speciation, extinction, and dispersal. Similar species richness disparities may be formed by differences in either the rate or timing of these processes (Figure 1). We examined whether the rate of diversification (speciation minus extinction) or the length of time allowed for diversification (time since colonization) best explained the global distribution of freshwater fish richness. Overall, our results suggest that time-for-speciation is the lead driver of latitudinal species richness disparities. We also show that diversification rates are highest among recently colonized basins, suggesting that richness differences among this set of basins are driven by diversification rates. Time-for-speciation and diversification rates are therefore both needed to explain diversity patterns overall.

### 4.1 Time best explains latitudinal richness patterns

We found a latitudinal trend in time-for-speciation associated with the K-Pg boundary (Figures 3, 5). While dominant tropical lineages tend to have Mesozoic origins, the dominant temperate lineages tend to have Cenozoic origins (Figure 3C; Figure 5). These patterns are exemplified by two major groups of fishes: Otophysi and Percomorpha (Figure 5; Extended Results 1). The crown of Otophysi (142 mya in the phylogeny of Rabosky et al., 2018) was most often reconstructed as widespread in the Neotropics, Indo-Malay and Afrotropics. The Neotropic Characiformes, Gymnotiformes, and some Siluriformes are descended from this initial colonization. Several lineages nested within Otophysi colonized the Nearctic and Palearctic independently. The Nearctic and Palearctic members of Cyprinidae most likely arrived from Southeast Asia near the K-Pg boundary or shortly after during the Eocene (Figure 5; Figure S5). These patterns are consistent with past studies (Cavender, 1998; Briggs, 2005; Chen, Lavoué, & Mayden, 2013), and also using the alternative phylogeny of Betancur-R et al. (2015; Extended Results 1, Figure S5). In percomorphs, the colonization of the tropics also pre-dated colonization of the extratropics (Figure 5; Figure S5). For example, in many stochastic maps colonization of the Neotropics pre-dated the crown of Ovalentaria (108 mya), the clade that includes the Cyprinodontiformes and Cichlidae (Extended Results 1; Figure S5). These two lineages each contain younger temperate members (Cavender, 1998). These results are consistent with a body of evidence from fossil and molecular data that the modern Neotropical biota was assembled over long time scales rather than from recent diversification (Hoorn et al., 2010; Albert & Reis, 2011; Antonelli et al., 2018; Albert, Tagliacollo, & Dagosta, 2020).

**FIGURE 5.**
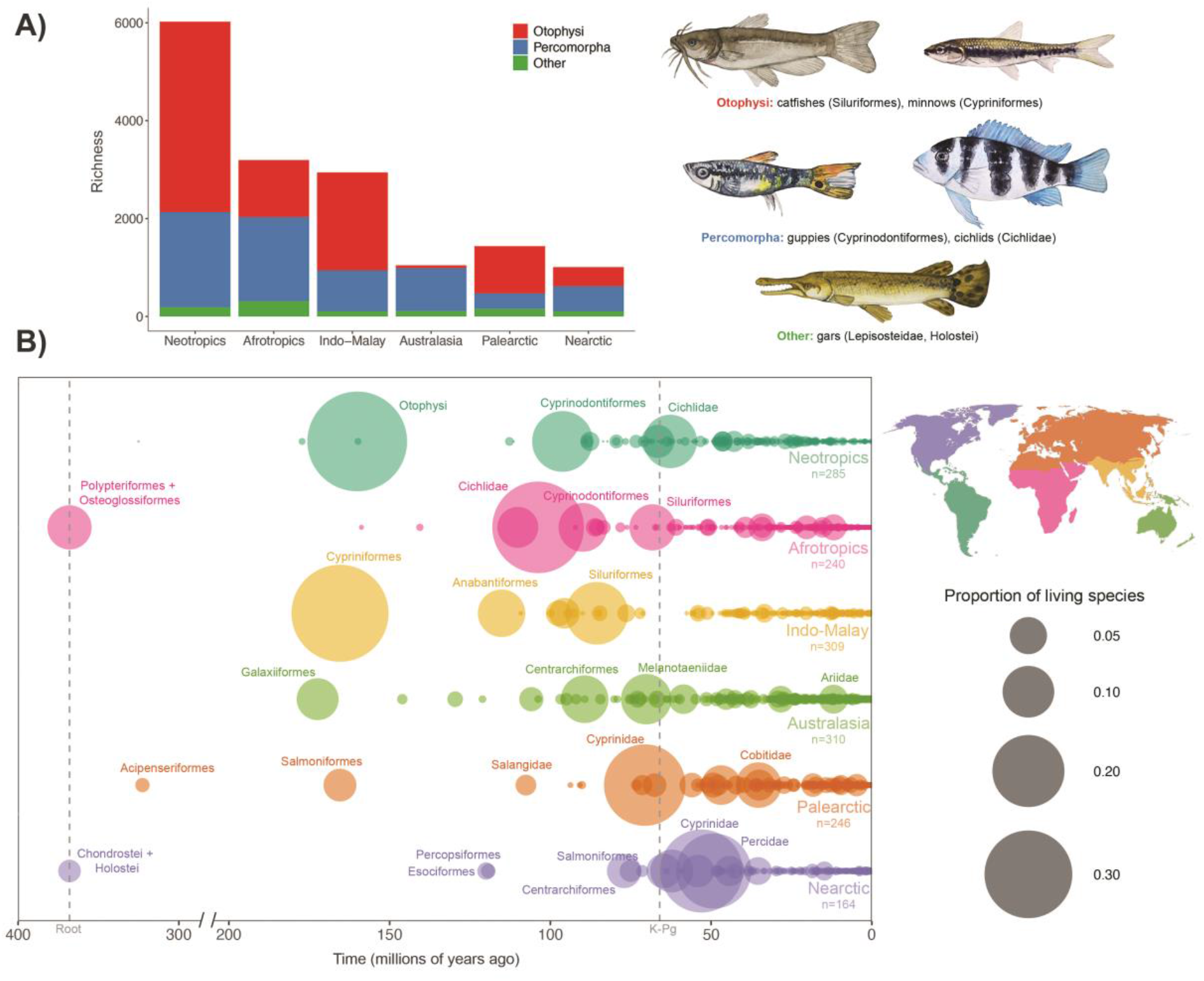
(**A**) Total regional species richness among three groups of ray-finned fishes: Otophysi (red), Percomorpha (blue), and all other groups combined (green). Richness estimates are from Tedesco et al. (2017). Exemplar members of each group are illustrated. Paintings by Kathryn Chenard. (**B**) Visualization of the relative contribution of colonization events though time. Each bubble represents a lineage that colonized independently; bubbles are scaled by the proportion of living species in the region descended from that event. The total number of colonizations of each region is noted. We used stochastic mapping (Dupin et al., 2016) and the phylogeny of Rabosky et al. (2018) to identify independently colonizing lineages and their living descendants. One example of a stochastic map is shown here; for variation among maps and with fossil data see Figure S5 and Extended Results 1. Mean and median times among stochastic maps were used in GAM analyses (Figures 2–4). Clade names represent the descendants of biogeographic events. Note that the colonization of a region can precede the crown of focal clades. See Figure S2 for an illustration of how this information was obtained from biogeographic reconstructions.

Richness also varies within the tropics, a pattern that can be attributed (in part) to colonization and time-for-speciation. While the Afrotropics contain ancient groups such as the Polypteriformes and Osteoglossiformes, communities today are dominated by the younger Cichlidae (Figure 5; Figure S5). Due to their numerical dominance, cichlids contribute more to the mean colonization times estimated for African basins than older groups (Figures 3, 5). While cichlids famously have fast diversification rates (McGee et al., 2020), they have not diversified over as long a period of time as the major Neotropical groups, and so the Afrotropics have comparatively lower richness today. The Australian tropics are unusual in that they were colonized many times, but no single lineage is dominant (Figure 5). The number of recent colonizations is consistent with greater ecological opportunity in the relatively depauperate Australian tropics (Betancur-R et al., 2012). The high endemism within Australian drainage basins suggests that the lack of dominant groups is related to the inability of fishes to expand across the continent (Unmack, 2001).

### 4.2 Why time drives diversity patterns

What does time-for-speciation imply about the differences between low and high latitudes? The two hypotheses given below are not mutually exclusive and might have a combined effect in driving diversity patterns. One hypothesis may be more relevant for the origin of the pattern (barriers to colonization) and the other for its maintenance (extinction).

#### 4.2.1 Barriers to extratropical colonization

Our results for freshwater fishes align with the tropical conservatism model (Wiens & Donoghue, 2004) and the out-of-the-tropics model (Jablonski et al., 2006). The overall pattern of colonization conforms to the expectations of both models; that is, older tropical clades (e.g. Otophysi) exported nested lineages to extratropical regions (e.g. Cyprinidae). A prediction of the tropical conservatism model is that movement out of the tropics is limited by niche conservatism, or the tendency for related taxa to share similar environmental tolerances (Wiens & Donoghue, 2004). Under this model, we would expect tropical-extratropical dispersal to preferentially occur during warm periods in Earth’s history. The out-of-the-tropics model implies instead that niches of tropical clades are more evolutionarily labile than extratropical groups (Tomašových & Jablonski, 2017). Although we did not examine climactic niches in this study, some observations suggest that niche conservatism may indeed underlie colonization patterns in freshwater fishes. Many tropical clades have few or no living species that reach temperate latitudes today, including Characiformes, Gymnotiformes, and Cichlidae. In addition, major radiations found in the Holarctic (such as Cypriniformes and Percidae) arrived during a period when the Earth was much warmer overall compared to the present day (Figure 5; Mannion, Upchurch, Benson, & Goswami, 2014). Future work may investigate whether colonization preceded niche evolution in temperate fishes (Folk et al., 2019).

The Earth was in a greenhouse period for much of the evolution of modern freshwater fishes, from the mid-Permian to the Neogene (∼272 to 23 mya; Mannion et al., 2014). Why did the Holarctic otophysans and percomorphs arrive so late, if temperature was not a barrier? In addition to the low dispersal capabilities of freshwater fishes, there may have been other barriers to colonization. Laurasia and Gondwana were separated by the Atlantic Ocean by the time Otophysi and Percomorpha originated (142.1 and 122.7 mya respectively; Rabosky et al., 2018). Laurasian landmasses were also flooded by epicontinental seas to a greater extent than Gondwanan landmasses during the Cretaceous, and these seas began to recede during the early Cenozoic (Ronov, 1994). The flooding of Laurasia and its separation from Gondwana may have delayed its colonization by freshwater fishes, especially those with poor salt tolerance.

#### 4.2.2 Extinction and stability of the tropics

It has been traditionally thought that Quaternary glaciation cycles were important for explaining the latitudinal diversity gradient, because lower latitudes were unaffected by glaciation (Bush, 1994; Mittelbach et al., 2007). In freshwater fishes, Quaternary climate cycles have left a signature on beta diversity (community turnover) at high latitudes, suggesting that high latitudes were re-colonized by a subset of species after glaciers receded (Leprieur et al., 2011). Nonetheless, our reconstructions imply that the latitudinal diversity gradient in freshwater fishes was already established by the Cretaceous (Figures 3C, 5). Again, we must look to older events to fully explain the latitudinal diversity gradient.

The Osteoglossiformes, Characiformes, Cyprinodontiformes, and Channidae all show fossil evidence of extinction in high latitudes during the Cenozoic (Lavoué, 2016; Capobianco & Friedman, 2018). Meseguer & Condamine (2020) pointed out that if the latitudinal diversity gradient was flatter during warm periods in Earth’s past (Mannion et al., 2014; Saupe et al., 2019), then temperate extinctions and range contractions must have played a role in the sharp gradient we see today. They suggested that this asymmetric extinction and dispersal model is an extension of the tropical conservatism hypothesis (Wiens & Donoghue, 2004): when warm climates became restricted to low latitudes, warm-adapted temperate lineages went extinct. Therefore, extinction, colonization and time-for-speciation can be related. Recent arrivals to the temperate zone have had limited time to replace diversity lost from extinction and range contraction (see also Miller & Wiens, 2017). Tropical groups were less affected by range contraction, and so were allowed to diversify over longer periods of time.

### 4.3 Relation to other hypotheses

While we focused on diversification rates and time in this manuscript, our results have implications for other potential influences on species richness.

#### 4.3.1 Ecological limits

The presence of ecological limits is often tested with the correlation between species richness and proxy variables such as productivity (e.g. Machac, 2020). Regions with low productivity are thought to have smaller carrying capacities, and diversification rates should slow more quickly than regions with higher carrying capacities (Rabosky, 2009). Our results are seemingly in conflict with this prediction. We found an inverse relationship between time and diversification rates: species-rich tropical basins had modest rates, while recently colonized but depauperate basins had the fastest rates (Figure 3, 4). Basins with fast rates tended to be found in arid regions (Figure 3; Smith et al., 2010). This inverse pattern is also seen in simulations (Hurlbert & Stegen, 2014) and empirically in marine fishes (Rabosky et al., 2018; Miller et al., 2018), marine invertebrates (O’Hara et al., 2019) and birds (Machac, 2020), and even has precedents in older literature (Briggs, 1966). These observations suggest that using environmental variables to test for ecological limits can give misleading results (Buckley et al., 2010). Studies that use a phylogenetic approach to trace the evolution of biogeographic ranges may have greater potential to reveal how ecological limits act on diversification and colonization (Betancur-R et al., 2012; Moen & Morlon, 2014).

Machac (2020) suggested that the ecological limits, diversification rates and time hypotheses could be integrated to explain why old tropical faunas have depressed diversification rates. As species richness increases, diversification rates slow but are not zero, so that richness continues to increase. Species-poor regions should have fast diversification rates due to ecological opportunity, but low richness due to the limited time allowed for diversification. Our results can add to this integrated view. Basins with mean colonization times from ∼12–40 mya had the fastest rates, but basins with mean times from ∼40–150 mya did not vary strongly in rates (Figure 4). Lineages in the second set of basins may be past the “exponential growth” phase (Rabosky, 2009). Once diversification rates begin to slow it will take more time to add new species, such that species richness among mature faunas may be better explained by time-for-speciation.

#### 4.3.2 Dispersal frequency

Biased dispersal rates can contribute to species richness disparities without invoking differences in diversification rates or time (Roy & Goldberg, 2007). We found that richness was strongly and positively related to the number of lineages in each basin representing independent colonization events. This is expected: a basin with one colonizing lineage can have one or more species, but a basin containing five independently colonizing lineages cannot have fewer than five species; and so on.

We may expect a positive relationship between time and the number of colonizations: the longer an area has been available and suitable for freshwater fishes, the more colonists it should accrue. Surprisingly, we found a weak negative relationship between the number of colonizations and mean colonization time (Figure S4). We think this is due to the asymmetric contribution of individual colonization events to richness. Much freshwater fish diversity is derived from only a few lineages. The first arrival of the Neotropics by the Otophysi accounts for at least ∼40% of present-day richness in the region (Figure 5). Biogeographic patterns in freshwater fishes therefore contrast with those for marine fishes, where colonization frequency is a major driver of regional richness (Miller et al., 2018). This suggests that in freshwater, where dispersal capabilities are low, high species richness is more easily achieved though in-situ speciation (Figure 1) than from high dispersal rates. If colonization rates are high, but have only been high recently, then the corresponding effect on richness may be limited especially if most new colonists become locally extinct over tens of millions of years.

#### 4.3.3 Area

Area has traditionally been considered alongside time as an important predictor of species richness (Willis, 1922). We found that surface area of drainage basins was related to species richness (Figure S1) in congruence with Oberdorff et al. (1995). However, latitude was a poor predictor of basin area, and time-for-speciation was more closely related to latitude than area. This suggests that latitudinal trends in time-for-speciation are not a response to an area-latitude relationship. One possibility is that past area is a better predictor of latitudinal diversity patterns than present-day area (Fine & Ree, 2006). Continental flooding would have reduced the area accessible to freshwater fishes in the Holarctic (Ronov, 1994), which may have delayed colonization. In conjunction with time, this could help explain why some Holarctic basins are less species rich than expected given their size today (e.g. the Mississippi; Figure 3A). Speciation takes time to complete, and long periods of time might be needed to build richness even if the area is large enough to support many species (Li & Wiens, 2019).

### 4.4 Caveats and sources of error

#### 4.4.1 Sampling and divergence times

Fishes are less densely sampled for genetic information than other vertebrates (Jetz et al., 2012), and knowledge of fish communities varies by region (Tedesco et al., 2017). We avoided inferring trends in diversification rates through time, which are sensitive to sampling (Blackburn, Giribet, Soltis, & Stanley, 2019) and model identifiability (Louca & Pennell, 2020). Note however that the DR estimates used here were inferred from a phylogeny with missing species imputed with taxonomic constraints (Rabosky et al., 2018). Our results based on DR estimates were similar to those using BAMM rates inferred from the tree with genetic data only (Tables S3–S6), suggesting that variation in the magnitude of diversification rates among fishes is not likely to be driven by sampling biases.

We believe that the temporal patterns of colonization inferred here should be robust. While the phylogeny used contains ∼36% of living actinopterygian species, it includes 90.2% of families and 100% of orders (Rabosky et al., 2018). This degree of sampling is sufficient to capture the age of higher taxa in the tree (Sanderson, 1996) which are more relevant to our time-for-speciation results than the most recent nodes. We compared colonization times estimated from this phylogeny and an alternative with greatly reduced sampling and different divergence time estimates (Betancur-R et al., 2015). The order of events was still generally common to both trees, with clades originating in the tropics and exporting nested lineages to the temperate zone (Extended Results 1; Figure S5). Sampling and divergence time estimates will both improve with more study, but as long as these nested relationships are preserved then tropical colonizations will still tend to be older than temperate colonizations (Cavender, 1998; Chen et al., 2013).

#### 4.4.2 Range estimates

The methods used here estimate ancestral ranges using the relationships and present-day distribution of living species. Range contractions from the temperate zone are known from the Cenozoic fossil record of fishes (Cavender, 1998; Lavoué, 2016; Capobianco & Friedman, 2018). Whether or not our results will be biased by the lack of recent fossils in our tree depends on the phylogenetic relationships between the missing fossils and living taxa. If the missing temperate fossils represent independent colonizations with no living descendants, then they will have little bearing on the ancestral reconstructions among living groups. If these fossils are instead placed near the stem branch of living temperate groups, then the timing of temperate colonizations could be underestimated from extant data. Even so, there is strong evidence that modern tropical lineages have been present since the Mesozoic, and temperate lineages tend to be nested within tropical ones (Figure 5). Therefore, this potential underestimation should not overturn the general trend for tropical colonizations to precede temperate colonizations. We predict that the continued integration of fossils and molecular phylogenies will refine colonization time estimates, but should not overturn our conclusions about the role of time on the latitudinal diversity gradient (Extended Results 1 and 2, Figure S5). As mentioned, these range contractions may actually support, not contradict, the importance of time-for-speciation (Miller & Wiens, 2017; Meseguer & Condamine, 2020).

## 5 CONCLUSIONS

We show that the latitudinal diversity gradient in freshwater fishes is primarily driven by earlier colonization of low-latitude regions, extending the timeline of diversification in the tropics compared to higher latitudes. More broadly, our results suggest that the most likely path to building very high species richness is through diversifying over long periods of time rather than diversifying very quickly. A remaining question is whether we observe younger temperate colonizations because the temperate zone has been harder to colonize, because new colonizations are replacing older ones that went extinct, or other scenarios. The time-for-speciation effect may manifest through colonization opportunity, niche conservatism, ecological limits, environmental stability, and/or past extinction (Stephens & Wiens, 2003; Wiens & Donoghue, 2004; Miller & Wiens, 2017; Pontarp & Wiens, 2017). A priority for future research is determining which of these factors are most relevant for generating the latitudinal diversity gradient, how they interact, and why they vary in space and time.

## ACKNOWLEDGEMENTS

We thank John J. Wiens and Andrew Zaffos for discussion and support. This manuscript was improved with comments from Adam Tomašových, Ricardo Betancur-R, and an anonymous reviewer. Kathryn Chenard painted the fishes in Figure 5. Elizabeth Miller was supported by an NSF Graduate Research Fellowship (DGE-1143953) and NSF Postdoctoral Fellowship (DBI-1906574).

## DATA AVAILABILITY STATEMENT

The data utilized in this study are available at https://figshare.com/s/552e7549303de8f2a823.

## Supplementary material associated with this manuscript

### Supplementary text

1. Extended Methods
2. Extended Results 1: Comparing colonization time estimates between two phylogenies, one with fossil taxa
3. Extended Results 2: Effect of excluding early diverging lineages on diversification rate and colonization time estimates of basins

### Supplementary tables

1. Table S1: Constraints on dispersal used in stratified DEC model
2. Table S2: Regions assigned to fossil taxa with references
3. Table S3: Change in richness, diversification rates, time-for-speciation and surface area with latitude and longitude
4. Table S4: Effect of time-for-speciation and diversification rates on species richness
5. Table S5: Effect of time-for-speciation and diversification rates on species richness while controlling for the species-area scaling
6. Table S6: Relationship between diversification rates and colonization times
7. Table S7: Influence of area on trends in diversification rates and time-for-speciation
8. Table S8: Relationship between number of colonizations and richness, latitude, time and diversification rates

### Supplementary figures

1. Figure S1: The relationship between basin surface area and richness
2. Figure S2: Method to illustrate complex colonization-richness temporal dynamics
3. Figure S3: Relationship between richness and diversification rates or time
4. Figure S4: Number of colonizations represented in each basin, and relationship to richness, latitude, time and diversification rates
5. Figure S5: Comparing colonization time estimates between alternative phylogenies

