## Supporting Information for "Evolutionary time best explains the latitudinal diversity gradient of living freshwater fish diversity"

### SUPPLEMENTARY INFORMATION

*Included in this document:*

Supplementary text:
1. **Extended Methods**
2. **Extended Results 1:** Comparing colonization time estimates between two phylogenies, one with fossil taxa
3. **Extended Results 2:** Effect of excluding early diverging lineages on diversification rate and colonization time estimates of basins

Supplementary tables:
1. **Table S1:** Constraints on dispersal used in stratified DEC model
2. **Table S2:** Regions assigned to fossil taxa with references
3. **Table S3:** Change in richness, diversification rates, time-for-speciation and surface area with latitude and longitude
4. **Table S4:** Effect of time-for-speciation and diversification rates on species richness
5. **Table S5:** Effect of time-for-speciation and diversification rates on species richness while controlling for the species-area scaling
6. **Table S6:** Relationship between diversification rates and colonization times
7. **Table S7:** Influence of area on trends in diversification rates and time-for-speciation
8. **Table S8:** Relationship between number of colonizations and richness, latitude, time and diversification rates

Supplementary figures:
1. **Figure S1:** The relationship between basin surface area and richness
2. **Figure S2:** Method to illustrate complex colonization-richness temporal dynamics
3. **Figure S3:** Relationship between richness and diversification rates or time
4. **Figure S4:** Number of colonizations represented in each basin, and relationship to richness, latitude, time and diversification rates
5. **Figure S5:** Comparing colonization time estimates between alternative phylogenies

References

*Included in FigShare repository* (doi: 10.6084/m9.figshare.8251394):

1. **Table A1:** Presence Absence Matrix (PAM) summarizing the distribution of 14,947 species across 3,119 basins
2. **Table A2:** Basin-specific mean rates of diversification based on BAMM and DR
3. **Table A3:** Species-specific mean colonization times derived from ancestral area reconstruction analyses.
4. **Table A4:** Basin-specific mean and median colonization times
5. **Table A5:** Basin-specific mean diversification rates, teleosts only
6. **Table A6:** Basin-specific mean colonization times, teleosts only

### Extended Methods

Additional details of our data and analyses are reported here.

***Estimating diversification rates for each freshwater drainage basin:***

Species occurrences within freshwater drainage basins were retrieved from Tedesco et al. (2017). This database can be downloaded from the online repository associated with this reference (https://doi.org/10.6084/m9.figshare.c.3739145). The original database includes occurrence records for 14,953 fish species across 3,119 basins found within seven biogeographic realms (Neotropics, Afrotropics, Indo-Malay, Nearctic, Palearctic, Australasia, and Oceania. We excluded occurrences marked in this database as non-native or questionable, and ultimately analyzed an occurrence dataset including 14,947 species.

Major freshwater fish clades are generally well sampled in the Tedesco et al. (2017) database. The database includes occurrence records for 93% of species in Characiformes, 93% of Gymnotiformes, 84% of Siluriformes, 71% of Cypriniformes, 79% of Osteoglossiformes, 61% of Anabantiformes, 75% of Melanotaeniidae, 73% of Cyprinodontiformes, 88% of Percidae, and 86% of Cichlidae (species counts were based on those from Rabosky et al., 2018).

We estimated the Presence-Absence Matrix ("PAM" hereafter; Gotelli, 2000; Arita, Christen, Rodríguez, & Soberón, 2008) of species occurrence across drainages (Table A1 in our FigShare repository). The PAM is a binary matrix summarizing in 14,947 rows (species) and 3,119 columns (drainage basins), and the occurrence of each species (either 1 or 0) within each basin. This PAM was later used to calculate the basin-specific rates of diversification.

Diversification rate estimates were based on time-calibrated molecular phylogenies constructed by Rabosky et al. (2018; also retrievable from https://fishtreeoflife.org). Diversification rates were estimated for each basin using two different approaches. Mean rates for each basin can be found in Table A2 in FigShare.

**(i) BAMM tips (six alternative rate estimates per basin)**: We analyzed the posterior distributions of six BAMM (Rabosky, 2014) runs published by Rabosky et al. (2018). BAMM output was available from the Dryad directory associated with the study. We analyzed output from three independent runs under a time-constant model of diversification and three independent runs under a time-variable model. All BAMM analyses used the same topology: the maximum likelihood phylogeny including species with genetic data only (11,686 species). BAMM event data were loaded into R using the getEventData function implemented in the 'BAMMtools' R package (Rabosky et al., 2014). Using rates from each of six BAMM runs, we calculated the mean net diversification rate (lambda minus mu) of each basin as the mean rate for species marked as present (i.e. rows with 1 in PAM) in each basin.

**(ii) DR (one rate estimated per basin):** We also estimated diversification rates using the DR statistic (Jetz, Thomas, Joy, Hartmann, & Mooers, 2012). DR tip rates estimated by Rabosky et al. (2018) were retrieved from the Dryad package associated with the study. These DR values were calculated using phylogenies with all unsampled species grafted using taxonomic constraints. DR values represent the means across a sample of 100 grafted phylogenies. Using the PAM for drainage basins, we estimated the DR value for each basin as the mean DR across all the species present.

#### *Estimating colonization times within basins:*

To estimate the timing of colonization of major regions (and therefore the amount of time allowed for diversification since colonization; Stephens & Wiens, 2003), we fit the dispersal-extinction-cladogenesis model (DEC; Ree & Smith, 2008) using the R package 'BioGeoBEARS' v1.1 (Matzke, 2014). Additional details of these reconstructions are given here (see Methods, main text). To assign each species in the phylogeny to one or more regions of occurrence (11,638 species with genetic data to begin), we first used the cleaned occurrence dataset of 14,947 species from Tedesco et al. (2017). We used FishBase (Froese & Pauly, 2019) to assign biogeographic regions to 631 species in the phylogeny that were missing from Tedesco et al. (2017). We removed 117 species from the phylogeny that were duplicates (multiple subspecies of the same species), unresolved to species level, or had unclear biogeographic affinity.

The computation time of biogeographic models scales exponentially with the number of areas allowed in the reconstruction (Matzke, 2014). To improve computational feasibility, we excluded Oceania from the reconstructions, a region limited to basins in French Polynesia, because only 17 species were endemic to this region. These 17 species were then removed from the phylogeny, and the basins in French Polynesia were discarded in any downstream comparisons using colonization times (see below). This lowered the total number of regions to seven (Neotropics, Afrotropics, Indo-Malay, Palearctic, Nearctic, Australasia, and "restricted to marine habitats"). In addition, while the majority of species were restricted to one or two regions, a few species were cosmopolitan. We removed 5 species occurring in more than three of the remaining biogeographic regions. The maximum number of areas allowed for any single lineage was then set to three. These steps greatly improved the computational feasibility of fitting a complex biogeographic model on this large phylogeny, while discarding as little data as possible. After these changes to the phylogeny and occurrence dataset, we performed ancestral area reconstructions using the resulting phylogeny of 11,499 species.

Our time-stratified model applied constraints on dispersal between regions, in accordance with changing regional connectivity through time. For consistency with prior literature, we followed Toussaint, Bloom, & Short (2017)'s DEC analyses for freshwater beetles. That is, dispersal between adjacent regions was not constrained (i.e. the probability of dispersal was equal to 1); dispersal probability among regions separated by a small marine barrier was set to 0.75; dispersal probability among regions separated by another landmass was set to 0.50; and dispersal probability among regions separated by a large marine barrier was set to 0.25. The dispersal probability between marine (the seventh "restricted to marine" category) and any continental region was set to 0.05 at all times to reflect the difficulty of these habitat transitions. These rules were applied to six time periods, spanning the root to the tips (0–20, 20–40, 40–80, 80–150, 150–350, and 350–368 mya). See further justification and details in Table S1.

#### *Comparing colonization times between phylogenies*

To assess the effect of fossils on our inferred colonization time, we also performed ancestral area reconstructions on the phylogeny from Betancur-R, Ortí, & Pyron (2015). This phylogeny includes 240 fossil and 1,582 extant species of ray-finned fishes. Some additional details of biogeographic coding are noted here (see also Table S2). To assign living species in this tree to biogeographic regions, we matched species to occurrences in the Tedesco et al. (2017) database. Species not found in this database were coded using FishBase (Froese & Pauley, 2019). We used Table SB1 from Betancur-R et al. (2015) to identify fossil taxa from freshwater regions (versus marine fossils). Some species were listed in this table as having uncertain habitat affinities. We assigned these fossils to freshwater regions based on where they were found. This approach is congruent to ours using the Rabosky et al. (2018) phylogeny, where brackish and marine species were coded as present in freshwater regions if they had occurrences in freshwater drainage basins in the Tedesco et al. (2017) database. We excluded 1 fossil species from Antarctica (region not included in this study). We also excluded 1 living species that was cosmopolitan (found in all 6 regions) to limit the computational intensity of the analysis. The resulting phylogeny contained 90 freshwater and 149 "exclusively marine" fossils, as well as 1,581 extant taxa.

Analyses using BioGeoBEARS were performed identically to those using the Rabosky et al. (2018) phylogeny, with some exceptions noted here. One fossil species (*Semionotus*) was found in five regions. Rather than exclude this fossil, we set the maximum number of regions to 5 (instead of 3). We also slightly adjusted the boundaries of some time periods used for stratifying the tree and constraining dispersal. For example, instead of 40–80 mya we used 40.5–80.5 mya (see all adjustments in Table S1). This was because the stratification fails when these boundaries are identical to the age of some nodes in the tree.

#### *Testing the relationships between diversification rates, time-for-speciation and species richness across biogeographical realms:*

First, we used spatially-explicit Generalized Additive Models (GAMs) to examine how species richness, net diversification rates, time-for-speciation and surface area are related to latitude, longitude and the interaction between latitude and longitude. GAMs were fitted using GAM function in the 'mgcv' R package (R Core Team, 2019; Wood, 2011). We also used correlation analyses based on the Spearman test to estimate the strength and direction of the association between geographic position and each of the analyzed predictors. Correlations were fitted using the cor.test function in R ('states' package; R Core Team, 2019). We analyzed absolute values of latitude in models that did not assume the interaction with longitude. We log-transformed richness and diversification rates and used raw values of colonization times. We used a log-link for models analyzing richness as response. These results are in Table S3.

Second, we estimated the effect of time-for-speciation and diversification rates on species richness, in isolation and relative to each other (Tables S4, S5; Figure S3). We estimated the relative contributions of time-for-speciation versus diversification rates for explaining spatial diversity patterns using four models ("null_model", "div_model", "time_model", and a full model). The amount of deviance explained by time-for-speciation was: (deviance(time_model)-deviance(full_model))/deviance(null_model). The amount of deviance explained by diversification rates was: (deviance(div_model)-deviance(full_model))/deviance(null_model). Note that the larger the difference in deviance for the sub-model relative to the full model, the more important the excluded predictor is (i.e. the absence of a given predictor implies a larger information loss relative to the full model). All nested models were fit by constraining estimated parameters from the full model using the sp parameter in the gam function. We also performed these analyses with the addition of surface area as a covariate (Table S5).

Third, we tested if diversification rates and colonization times are related to each other across basins. We analyzed spatially explicit GAMs and non-parametric correlation tests between colonization times (predictor) and diversification rates. These results are in Table S6.

Fourth, we used GAMs to examine whether latitudinal gradients in time-for-speciation and diversification rates are better explained by a co-variation between area and latitude. Below, we explain the procedure we follow for colonization times. First, we fit a GAM with area and latitude as predictors of colonization times. Next, we fit two additional models with colonization times being predicted independently by latitude or area. We estimated the fraction of deviance explained in the full model (area + latitude) versus each of the predictors (area or latitude) by comparing $R^2$ values of each model and the change in deviance after excluding each predictor from the full model. The same procedure was repeated to analyze the effects of area and latitude in driving spatial patterns in diversification rates. These results are in Table S7.

Finally, we tested whether the number of regional colonizations represented among species in each basin was related to basin richness, latitude, diversification rates, or colonization times. See Figure S2 for an illustration of how we identified colonizations. These results are in Table S8 and Figure S4.

**Extended Results 1.** Comparing colonization time estimates between two phylogenies, one with fossil taxa

The purpose of this section is threefold: (1) to assess how including fossils may influence our results; (2) to discuss the uncertainty surrounding some estimates of colonization time; and (3) to give more detail on the temporal biogeographic patterns seen among fishes.

The phylogeny from Rabosky et al. (2018; hereafter "RAB") contains 11,499 species, 422 families (90.2%) and 68 orders (100%) with geographic data. An earlier phylogeny from Betancur-R et al. (2015; hereafter "BET") is more sparsely sampled with 1,582 extant species, though it contains 378 families (77.1%) and 68 orders (100%). A major benefit of the BET phylogeny is that it also contains 240 fossil taxa, including marine stem members of clades that are now restricted to freshwater. Thus, it can be used to test if freshwater colonization times may be overestimated for some groups using the RAB phylogeny.

It is not possible to compare all colonization times due to the difference in sampling between the two trees. Instead, we compared the range of dates of important biogeographic events sampled from 100 simulations of biogeographic history ("biogeographic stochastic mapping") on either phylogeny. These events are the first arrivals of select clades to regions. We selected 15 clades to compare, chosen because they are early-diverging, or because they make up a large portion of the extant diversity of continental regions (Figure 5). Note that in many cases, the occupancy of the region predated the crown age of the clade (see Figures S2 and S5). Note also that while we discuss the range of dates sampled among simulations here, the mean and median dates among simulations were used in our GAM analyses (e.g. Figures 2–4). See Figure S5 for a visualization of dates described herein.

*Early-diverging lineages:* Fossil taxa in the BET tree helped to restrict the range of likely colonization times, and the mean dates tended to be much later in the BET tree compared to the RAB tree. **Due to these differences, we tested whether removing early diverging groups was likely to influence our results (Extended Results 2).**

A. **Polypteriformes in Afrotropics:** The ancestral range reconstructed at the root of the RAB phylogeny included the Afrotropics in all simulations. As such, the occupancy of the Afrotropics preceded the origin of the Polypteriformes (bichirs), at the root of all ray-finned fishes (~368 million years ago). There was great uncertainty in the time of colonization of the Afrotropics using the BET tree, due to fossil taxa near the root and the long stem branch subtending the Polypteriformes. The mean time of colonization among 100 simulations was 279.1 mya, much later than the RAB tree.

B. **Acipenseriformes in Palearctic:** Note that the Acipenseriformes here include the sturgeons and paddlefishes. There was great uncertainty in this event using the RAB tree, with estimates spanning the root to 56 mya (mean 284.4 myr). The estimate using the BET tree was restricted to ~252–140 mya (mean 188.2 myr) due to marine stem fossils breaking up the branch leading to the crown group.

C. **Acipenseriformes in Nearctic:** Similar to (B), there was great uncertainty in the timing of this event using the RAB tree, from root to ~68 mya (mean 347.9 myr). The estimate using the BET tree was restricted to ~242–100 mya (mean 183.6 myr).

D. **Osteoglossiformes in Afrotropics:** There was great uncertainty in the timing of arrival using the RAB tree compared to the BET tree, due to the lack of fossil taxa. However, both trees agreed that the occupancy of the Afrotropics most likely preceded the origin of the Osteoglossiformes. The difference in mean times between the two trees was not as great as for clades A–C (mean 234.1 myr for RAB and 195.7 myr for BET).

*Otophysan lineages:* In both phylogenies, the relationships among Otophysan orders are (Cypriniformes, (Gymnotiformes, (Characiformes, Siluriformes))). The relationships within the Cypriniformes are (Cyprinidae, (Catostomidae, loach families). The congruence of these relationships between trees means that the timing of biogeographic events is largely congruent as well. In both RAB and BET trees, the mean colonization times for the most diverse clades are ordered as follows from oldest to youngest: Neotropics, Indo-Malay, Palearctic, and Nearctic.

E. **Otophysi in Neotropics:** Both trees showed that the occupancy of the Neotropics most likely preceded the origin of the Otophysi. Most Neotropical Otophysans (Characiformes, Gymnotiformes, and some Siluriformes) are descended from this initial event (Figure 5). Using the RAB phylogeny, there was a non-zero but unlikely probability of this event being exceptionally old (e.g. if the root of the tree was reconstructed in the Neotropics). Still, the mean date among 100 simulations was actually younger in the RAB tree (160.3 mya) than the BET tree (195.1 mya).

F. **Cypriniformes in Indo-Malay:** The following features are shared by both trees. The Cypriniformes were the first Otophysan lineage to occupy the Indo-Malay region and are also the most diverse lineage there today. Whether this colonization event occurred before or after the origin of the Otophysi varies among simulations. The mean dates are similar between trees (137.8 myr, RAB and 145.9 myr, BET). The Siluriformes dispersed to the Indo-Malay region independently and much later than the Cypriniformes (Figure 5).

G. **Cyprinidae in Palearctic:** Otophysan lineages colonized the Palearctic several times independently. The most diverse of these lineages is the Palearctic cyprinids. While this lineage was not the first within Otophysi to colonize the Palearctic (this was the catostomid *Myxocyprinus asiaticus*), it has the largest influence on our main results because it is numerically dominant within drainage basins. Using the RAB tree, this lineage most likely arrived in the Palearctic around the K-Pg boundary (mean 64.7 mya). There is considerable uncertainty using the BET tree, and the mean date is much earlier (range ~224–48 mya; mean 124 mya). We believe this is due to sampling differences between the trees. Fewer Palearctic otophysans are sampled in the BET tree; for example, the species of *Myxocyprinus* that likely colonized the Palearctic independently was not sampled in the BET tree. This means that the branch subtending the Palearctic cyprinids is longer and the time of colonization appears older and more uncertain.

H. **Cyprinidae in Nearctic:** This event shares many features with event G. Otophysans have colonized the Nearctic several times, and the oldest event is associated with Catostomidae. The most numerically dominant lineage is the Nearctic cyprinids, which we focus on here. This lineage colonized the Nearctic after the K-Pg boundary in all simulations using the RAB tree (mean 49.1 mya). There is greater uncertainty using the BET tree, and an older inferred time (range 216–28 mya, mean 85.9 mya). Again, this may be attributed to sampling differences between the trees.

*Percomorph lineages:* Estimates of colonization times among percomorph clades are generally congruent between the RAB and BET trees, and the uncertainty in these estimates is smaller than for older fish groups (Figure S5). Of 7 clades discussed here, 5 are in the series Ovalentaria.

**Cyprinodontiformes in Neotropics (I) and Afrotropics (J); Cichlidae in Neotropics (K) and Afrotropics (L):** The biogeographic patterns of these groups are related, so we discuss them together. In ~¾ of simulations using the RAB tree (76 of 100), a colonization of the Neotropics occurred at the crown of Ovalentaria or earlier. As a consequence, in most simulations the Cyprinodontiformes and Cichlidae are descended from the same colonization of the Neotropics. These two groups colonized the Afrotropics independently. Colonization of the Afrotropics occurred just prior to the crown of Cichlidae in all simulations, and within Cyprinodontiformes in most (71 of 100) simulations. Therefore, using the RAB tree the mean colonization time of the Neotropics was almost identical for the Cyprinodontiformes and Cichlidae (107.2 and 107.6 mya respectively), and preceded the mean colonization time of the Afrotropics by each group (Cyprinodontiformes: 90.1 mya; Cichlidae: 85.1 mya).

The reconstructions using the BET tree showed a reversal of this pattern, with the colonization of the Afrotropics generally preceding that of the Neotropics (Figure S5). In 76/100 simulations, a colonization of the Afrotropics occurred at the crown of Ovalentaria or earlier. The Cyprinodontiformes and Cichlidae then colonized the Neotropics independently (in 93/100 simulations). This difference between trees is likely due to the overall denser sampling of the RAB tree. For example, if more Neotropical members of Ovalentaria are sampled, an earlier reconstruction of the Neotropics may fit the data better depending on the distribution of these species within the series.

**Anabantiformes in Indo-Malay (M), Melanotaeniidae in Australasia (N), and Percidae in Nearctic (O):** For these three clades, the mean colonization time and range among simulations were very similar between the RAB and BET trees, so we do not discuss them further.

**Extended results 2**: The effect of excluding early diverging lineages on rate and time means

Our biogeographic reconstructions using a phylogeny with fossil taxa (Betancur-R et al., 2015) showed that colonization times associated with early-diverging clades were likely to be greatly overestimated using the Rabosky et al. (2018) phylogeny (Extended Results 1; Figure S5). To assess the impact of these groups on our results, we excluded the 48 species from the four living non-teleost clades: Polypteriformes, Acipenseriformes, Lepisosteiformes, and Amiiformes. We then re-estimated basin-specific rates of net diversification and mean colonization times.

Excluding non-teleosts altered rate and time estimates in 23% of drainage basins globally (725 of 3,119). Most of these basins (76%) were located in the Nearctic (213 basins) or Palearctic (338 basins).

**Rates of diversification (based on 12k_tc1; Table A6).** The exclusion of non-teleosts did not significantly change diversification rate estimates globally (t-test between dataset with and without non-teleosts: t = -0.167, df = 6101.2, P-value = 0.434). Mean diversification rates in the original dataset were on average 0.24% slower than in the new dataset excluding early diverging clades (sd=1.22%). Therefore, the exclusion of these early diverging clades had a small positive (but non-significant) effect on overall rate estimates.

**Mean colonization times (Table A7).** The exclusion of non-teleosts did not significantly affect mean colonization times across all basins (t-test between dataset with and without non-teleosts: t= 0.802, df = 6101.2, P-value = 0.43). Mean colonization times in the original dataset were on average 1.80% older compared to the dataset excluding early diverging clades (sd=7.30%). Therefore, the exclusion of these early diverging clades had a small negative (but non-significant) effect on mean colonization times overall. This small effect would increase the gap between low and high-latitude colonization times, supporting our conclusions based on all actinopterygians.

To sum, the exclusion of non-teleosts had little effect on mean diversification rates and time-for-speciation of basins overall. Therefore, excluding non-teleosts would not overturn our conclusions on the importance of time versus diversification rates for explaining latitudinal patterns in freshwater fish diversity, and may even further support our conclusions. Note that non-teleosts seem to have had little effect on basin means because they are species-poor (comprising only 48 species in four orders today). Our results are primarily being driven by numerically dominant lineages (Figure 5).

**Table S1:** Constraints on dispersal between regions by time period used in stratified DEC model fitting (Ree & Smith, 2008; Matzke, 2014). We constrained dispersal probabilities with time in concordance with tectonic activity and changing connectivity among regions. For consistency with prior literature, we followed Toussaint et al. (2017)'s DEC analyses for freshwater beetles. That is, dispersal between adjacent regions was not constrained; dispersal probability among regions separated by a small marine barrier was set to 0.75; dispersal probability among regions separated by another landmass was set to 0.50; and dispersal probability among regions separated by a large marine barrier was set to 0.25. The dispersal probability between marine and freshwater habitats was set to 0.05 at all times to reflect the difficulty of habitat transitions. Note that the boundaries of some time periods were adjusted when using the Betancur-R et al. (2015) phylogeny, because the stratification fails when these boundaries are identical to the age of some nodes in the tree.

**0-20 mya**: Nearctic and Neotropic regions connected by Isthmus of Panama; Tethys Ocean closed connecting Africa and Europe; Nearctic and Europe intermittently connected

| | Neotropics | Palearctic | Nearctic | Afrotropics | Australasia | Indo-Malay | Marine |
|---|---|---|---|---|---|---|---|
| Neotropics | - | 0.25 | 1 | 0.25 | 0.25 | 0.25 | 0.05 |
| Palearctic | 0.25 | - | 0.75 | 1 | 0.5 | 1 | 0.05 |
| Nearctic | 1 | 0.75 | - | 0.25 | 0.25 | 0.25 | 0.05 |
| Afrotropics | 0.25 | 1 | 0.25 | - | 0.25 | 0.5 | 0.05 |
| Australasia | 0.25 | 0.5 | 0.25 | 0.25 | - | 1 | 0.05 |
| Indo-Malay | 0.25 | 1 | 0.25 | 0.5 | 1 | - | 0.05 |
| Marine | 0.05 | 0.05 | 0.05 | 0.05 | 0.05 | 0.05 | - |

**20-40* mya**: India closer to Africa via Arabian Peninsula; Australia approaching Indo-Malay; Isthmus of Panama not connected; Africa and Europe separated by Tethys Ocean; Europe and Nearctic connected through Beringia
*adjusted to 40.5 when using the phylogeny from Betancur-R et al. (2015)

| | Neotropics | Palearctic | Nearctic | Afrotropics | Australasia | Indo-Malay | Marine |
|---|---|---|---|---|---|---|---|
| Neotropics | - | 0.25 | 0.75 | 0.25 | 0.25 | 0.25 | 0.05 |
| Palearctic | 0.25 | - | 1 | 0.75 | 0.25 | 1 | 0.05 |
| Nearctic | 0.75 | 1 | - | 0.25 | 0.25 | 0.25 | 0.05 |
| Afrotropics | 0.25 | 0.75 | 0.25 | - | 0.25 | 0.75 | 0.05 |
| Australasia | 0.25 | 0.25 | 0.25 | 0.25 | - | 0.25 | 0.05 |
| Indo-Malay | 0.25 | 1 | 0.25 | 0.75 | 0.25 | - | 0.05 |
| Marine | 0.05 | 0.05 | 0.05 | 0.05 | 0.05 | 0.05 | - |

**40*-80* mya**: South America and Africa are separated; Australia still connected to Antarctica
*adjusted to 40.5–80.5 when using the phylogeny from Betancur-R et al. (2015)

|  | Neotropics | Palearctic | Nearctic | Afrotropics | Australasia | Indo-Malay | Marine |
|---|---|---|---|---|---|---|---|
| Neotropics | - | 0.25 | 0.75 | 0.75 | 0.5 | 0.25 | 0.05 |
| Palearctic | 0.25 | - | 1 | 0.75 | 0.25 | 1 | 0.05 |
| Nearctic | 0.75 | 1 | - | 0.25 | 0.25 | 0.25 | 0.05 |
| Afrotropics | 0.75 | 0.75 | 0.25 | - | 0.25 | 0.75 | 0.05 |
| Australasia | 0.5 | 0.25 | 0.25 | 0.25 | - | 0.25 | 0.05 |
| Indo-Malay | 0.25 | 1 | 0.25 | 0.75 | 0.25 | - | 0.05 |
| Marine | 0.05 | 0.05 | 0.05 | 0.05 | 0.05 | 0.05 | - |

**80*-150* mya**: South America and Africa connected; India and Palearctic connected to Africa by land
*adjusted to 80.5–150.5 when using the phylogeny from Betancur-R et al. (2015)

|  | Neotropics | Palearctic | Nearctic | Afrotropics | Australasia | Indo-Malay | Marine |
|---|---|---|---|---|---|---|---|
| Neotropics | - | 0.5 | 0.75 | 1 | 0.5 | 0.5 | 0.05 |
| Palearctic | 0.5 | - | 1 | 1 | 0.25 | 1 | 0.05 |
| Nearctic | 0.75 | 1 | - | 0.25 | 0.25 | 0.25 | 0.05 |
| Afrotropics | 1 | 1 | 0.25 | - | 0.25 | 0.75 | 0.05 |
| Australasia | 0.5 | 0.25 | 0.25 | 0.25 | - | 0.75 | 0.05 |
| Indo-Malay | 0.5 | 1 | 0.25 | 0.75 | 0.75 | - | 0.05 |
| Marine | 0.05 | 0.05 | 0.05 | 0.05 | 0.05 | 0.05 | - |

**150*-350 mya:** Pangaea was intact
*adjusted to 150.5 when using the phylogeny from Betancur-R et al. (2015)

|  | Neotropics | Palearctic | Nearctic | Afrotropics | Australasia | Indo-Malay | Marine |
|---|---|---|---|---|---|---|---|
| Neotropics | - | 0.5 | 1 | 1 | 0.5 | 0.5 | 0.05 |
| Palearctic | 0.5 | - | 1 | 0.5 | 0.25 | 1 | 0.05 |
| Nearctic | 1 | 1 | - | 1 | 0.25 | 0.25 | 0.05 |
| Afrotropics | 1 | 0.5 | 1 | - | 0.5 | 1 | 0.05 |
| Australasia | 0.5 | 0.25 | 0.25 | 0.5 | - | 1 | 0.05 |
| Indo-Malay | 0.5 | 1 | 0.25 | 1 | 1 | - | 0.05 |
| Marine | 0.05 | 0.05 | 0.05 | 0.05 | 0.05 | 0.05 | - |

**350 mya-root of phylogeny*:** Rheic ocean separates Euramerica and Gondwana
*Root is dated at 368 mya for Rabosky et al. (2018), and 403 mya for Betancur-R et al. (2015)

|  | Neotropics | Palearctic | Nearctic | Afrotropics | Australasia | Indo-Malay | Marine |
|---|---|---|---|---|---|---|---|

| | | | | | | | |
|---|---|---|---|---|---|---|---|
| Neotropics | - | 0.25 | 0.75 | 1 | 0.5 | 0.5 | 0.05 |
| Palearctic | 0.25 | - | 1 | 0.25 | 0.25 | 1 | 0.05 |
| Nearctic | 0.75 | 1 | - | 0.75 | 0.25 | 0.25 | 0.05 |
| Afrotropics | 1 | 0.25 | 0.75 | - | 0.5 | 1 | 0.05 |
| Australasia | 0.5 | 0.25 | 0.25 | 0.5 | - | 1 | 0.05 |
| Indo-Malay | 0.5 | 1 | 0.25 | 1 | 1 | - | 0.05 |
| Marine | 0.05 | 0.05 | 0.05 | 0.05 | 0.05 | 0.05 | - |

**Table S2:** Regional coding for 90 fossil taxa in the phylogeny of Betancur-R et al. (2015).

| Taxon | Region(s) | References |
|---|---|---|
| *Aethalionopsis* | Palearctic | Grande & Poyato-Ariza, 1999; Cavin, 2017 |
| *Amia hesperia* | Nearctic | Cavin, 2017 |
| *Amia pattersoni* | Nearctic | Cavin, 2017 |
| *Amia scutata* | Nearctic | Cavin, 2017 |
| *Amiopsis damoni* | Palearctic | Martín-Abad & Poyato-Ariza, 2017 |
| *Amiopsis dolloi* | Palearctic | Martín-Abad & Poyato-Ariza, 2017; Cavin, 2017 |
| *Amiopsis prisca* | Palearctic | Martín-Abad & Poyato-Ariza, 2017 |
| *Amiopsis woodwardi* | Palearctic | Martín-Abad & Poyato-Ariza, 2017 |
| *Amphiplaga* | Nearctic | Cavin, 2017 |
| *Beishanichthys* | Palearctic | Cavin, 2017 |
| *Calamopleurus cylindricus* | Neotropics | Gardiner, Maisey, and Littlewood, 1996 |
| *Calamopleurus mawsoni* | Neotropics | Gardiner et al., 1996; Cavin, 2017 |
| *Chauliopareion* | Afrotropics | Wilson & Murray, 2008; Xu & Chang, 2009; Cavin, 2017 |
| *Cheirolepis canadensis* | Nearctic | Arratia & Cloutier, 1995 |
| *Cheirolepis trailli* | Palearctic | Arratia & Cloutier, 1995 |
| *Cretophareodus* | Nearctic | Cavin, 2017 |
| *Cuneognathus gardineri* | Nearctic | Friedman & Blom, 2006 |
| *Cyclurus efremovi* | Palearctic | Cavin, 2017 |
| *Cyclurus fragosus* | Nearctic | Cavin, 2017 |
| *Cyclurus gurleyi* | Nearctic | Cavin, 2017 |
| *Cyclurus ignotus* | Palearctic | Cavin, 2017 |
| *Cyclurus kehreri* | Palearctic | Cavin, 2017 |
| *Cyclurus macrocephalus* | Palearctic | Cavin, 2017 |
| *Cyclurus oligocenicus* | Palearctic | Cavin, 2017 |
| *Cyclurus valenciennesi* | Palearctic | Cavin, 2017 |
| *Dastilbe* | Neotropics | Grande & Poyato-Ariza, 1999; Cavin, 2017 |
| *Dialipina* | Palearctic, Nearctic | Schultze & Cumbaa, 2001 |
| *Diplomystus* | Palearctic, Nearctic | Cavin, 2017 |
| *Ellimmichthys* | Neotropics, Palearctic, Afrotropics | Cavin, 2017; Martill et al., 2011 |
| *Eohiodon rosei* | Nearctic | Frickhinger, 1995; Wilson & Murray, 2008; Xu & Chang, 2009 |
| *Eohiodon woodruffi* | Nearctic | Frickhinger, 1995; Wilson & Murray, 2008; Xu & Chang, 2009 |
| *Eolates aquensis* | Palearctic | Cavin, 2017 |
| *Erismatopterus* | Nearctic | Cavin, 2017 |
| *Evenkia* | Palearctic | Cavin, 2017 |
| *Fukangichthys* | Palearctic | Cavin, 2017 |
| *Gordichthys* | Palearctic | Grande & Poyato-Ariza, 1999; Cavin, 2017 |

| | | |
|---|---|---|
| *Hiodon consteniorum* | Nearctic | Wilson & Murray, 2008 |
| *Howqualepis rostridens* | Australasia | Friedman & Blom, 2006 |
| *Ikechaoamia meridionalis* | Palearctic, Indo-Malay | Hsien-Ting & De-Zao, 1983; Cavin, 2017 |
| *Ikechaoamia orientalis* | Palearctic, Indo-Malay | Hsien-Ting & De-Zao, 1983; Cavin, 2017 |
| *Jiaohichthys* | Palearctic | Jiang-Yong, 1998 |
| *Jiuquanichthys liui* | Palearctic | Jiang-Yong, 1998 |
| *Kuyangichthys microdus* | Palearctic | Jiang-Yong, 1998 |
| *Lateopisciculus* | Nearctic | Cavin, 2017 |
| *Libotonius* | Nearctic | Cavin, 2017 |
| *Limnomis delaneyi* | Nearctic | Friedman & Blom, 2006 |
| *Lycoptera davidi* | Palearctic | Cavin, 2017; Wilson & Murray, 2008; Jiang-Yong, 1998 |
| *Macrepistius arenatus* | Nearctic | Schaeffer, 1960 |
| *Mahengichthys* | Afrotropics | Cavin, 2017 |
| *Maliamia gigas* | Afrotropics | Cavin, 2017 |
| *Mansfieldiscus sweeti* | Australasia | Garvey & Hasiotis, 2008 |
| *Massamoricthys* | Nearctic | Murray, 1995 |
| *Mcconichthys* | Nearctic | Cavin, 2017 |
| *Melvius chauliodous* | Nearctic | Cavin, 2017 |
| *Melvius thomasi* | Nearctic | Cavin, 2017 |
| *Musperia radiata* | Indo-Malay | Wilson & Murray, 2008; Xu & Chang, 2009 |
| *Nipponamia satoi* | Palearctic | Cavin, 2017 |
| *Notelops* | Neotropics | Arratia, 2008 |
| *Notogoneus* | Palearctic, Nearctic, Australasia | Grande & Poyato-Ariza, 1999; Cavin, 2017 |
| *Novagonatodus kasantsevae* | Australasia | Garvey & Hasiotis, 2008 |
| *Ostariostoma* | Nearctic | Wilson & Murray, 2008; Cavin, 2017 |
| *Palaeonotopterus* | Palearctic | Wilson & Murray, 2008; Xu & Chang, 2009; Cavin, 2017 |
| *Parachanos* | Afrotropics | Grande & Poyato-Ariza, 1999; Cavin, 2017 |
| *Paraclupea* | Palearctic | Cavin, 2017 |
| *Paralycoptera wui* | Palearctic, Indo-Malay | Xu & Chang, 2009; Cavin, 2017; Jiang-Yong, 1998 |
| *Peipiaosteus* | Palearctic | Grande & Bemis, 1996 |
| *Phaerodusichthys tavernei* | Neotropics | Wilson & Murray, 2008 |
| *Phareodus encaustus* | Nearctic | Wilson & Murray, 2008; Xu & Chang, 2009; Cavin, 2017 |
| *Phareodus queenslandicus* | Australasia | Wilson & Murray, 2008; Xu & Chang, 2009; Cavin, 2017 |
| *Phareodus testis* | Nearctic | Wilson & Murray, 2008; Xu & Chang, 2009 |

| | | |
|---|---|---|
| *Plesiolycoptera daqingensis* | Palearctic | Jiang-Yong, 1998 |
| *Protopsephurus liui* | Palearctic | Cavin, 2017 |
| *Psammorhynchus longipinnis* | Nearctic | Cavin, 2017 |
| *Pseudamiatus heintzi* | Nearctic | Poyato-Ariza & Martín-Abad, 2013 |
| *Rhacolepis* | Neotropics | Arratia, 2008 |
| *Rubiesichthys* | Palearctic | Grande & Poyato-Ariza, 1999; Cavin, 2017 |
| *Santanaclupea* | Neotropics | Maisey, 1993 |
| *Scanilepis* | Palearctic | Ørvig, 1978 |
| *Semionotus* | Neotropics, Palearctic, Nearctic, Afrotropics, Australasia | Frickhinger, 1995 |
| *Sinamia zdanskyi* | Palearctic | Hsien-Ting & De-Zao, 1983; Gardiner et al., 1996; Cavin, 2017 |
| *Sinoglossus lushanensis* | Palearctic | Jiang-Yong, 1998 |
| *Stegotrachelus finlayi* | Palearctic | Friedman & Blom, 2006 |
| *Tanolepis ningjiagouensis* | Palearctic, Indo-Malay | Xu & Chang, 2009 |
| *Tharrias* | Neotropics | Grande & Poyato-Ariza, 1999 |
| *Thaumaturus* | Palearctic | Wilson & Murray, 2008; Cavin, 2017 |
| *Trichophanes* | Nearctic | Cavin, 2017 |
| *Vidalamia catalunica* | Palearctic | Cavin, 2017 |
| *Vinctifer* | Neotropics, Afrotropics | Cavin, 2017 |
| *Xixiaichthys* | Palearctic | Jiang-Yong, 2004 |
| *Yanbiania wangqingica* | Palearctic | Jiang-Yong, 1998 |

**Table S3.** Summary of results from Generalized Additive Models and Spearman correlation analyses testing the relationship between richness, basin area, diversification rates, and colonization times across latitude, longitude, and both latitude and longitude. Here we show the $R^2$ value for each of the fitted regression models, along with the associated P-value. Additionally, we also show both rho and P values based on non-parametric correlation analyses. See Figure S1B for the relationship between surface area and latitude.

| **Response variable** | | **Lat($R^2$)** | **lat(P)** | **lat(rho)** | **lat(P-rho)** | **long($R^2$)** | **long(P)** | **long+lat($R^2$)** | **long+lat(P)** |
|---|---|---|---|---|---|---|---|---|---|
| Richness | | 0.092 | <0.0001 | -0.273 | <0.0001 | 0.101 | <0.0001 | 0.257 | <0.0001 |
| Diversification | BAMM_tc1 | 0.207 | <0.0001 | -0.118 | <0.0001 | 0.034 | <0.0001 | 0.372 | <0.0001 |
| | BAMM_tc2 | 0.211 | <0.0001 | -0.118 | <0.0001 | 0.033 | <0.0001 | 0.375 | <0.0001 |
| | BAMM_tc3 | 0.210 | <0.0001 | -0.117 | <0.0001 | 0.033 | <0.0001 | 0.374 | <0.0001 |
| | BAMM_tv1 | 0.207 | <0.0001 | -0.121 | <0.0001 | 0.034 | <0.0001 | 0.369 | <0.0001 |
| | BAMM_tv2 | 0.205 | <0.0001 | -0.122 | <0.0001 | 0.033 | <0.0001 | 0.369 | <0.0001 |
| | BAMM_tv3 | 0.212 | <0.0001 | -0.111 | <0.0001 | 0.032 | <0.0001 | 0.371 | <0.0001 |
| | DR | 0.127 | <0.0001 | 0.334 | <0.0001 | 0.238 | <0.0001 | 0.390 | <0.0001 |
| Colonization | Mean | 0.271 | <0.0001 | -0.232 | <0.0001 | 0.116 | <0.0001 | 0.621 | <0.0001 |
| | Median | 0.253 | <0.0001 | -0.289 | <0.0001 | 0.143 | <0.0001 | 0.581 | <0.0001 |
| Surface area | | 0.054 | <0.0001 | -0.019 | 0.0233 | 0.085 | <0.0001 | 0.293 | <0.0001 |

**Table S4.** Results for spatially explicit Generalized Additive Models testing the relationship between richness, mean colonization times and diversification rates. Here we show the estimated $R^2$, P-value and deviance for four types of GAMs (boldfaced below). For bivariate models, we indicate the deviance explained by each predictor (dev.x%) along with the ratio of this index between variables (e.g. Time(%) ÷Div(%)). The larger this ratio is, the greater information lost compared to the full model by excluding time relative to diversification rates. Results are presented for 7 estimates of diversification rates and two different values of colonization times.

**Null model (Richness ~ 1):**

| $R^2$ | P | Deviance |
|---|---|---|
| 0.257 | <0.0001 | 2672.931 |

**Div model (Richness~Div):**

| Div rate estimate | $R^2$ | P | Deviance |
|---|---|---|---|
| BAMM_tc1 | 0.346 | <0.0001 | 2131.558 |
| BAMM_tc2 | 0.339 | <0.0001 | 2152.685 |
| BAMM_tc3 | 0.338 | <0.0001 | 2155.699 |
| BAMM_tv1 | 0.346 | <0.0001 | 2129.899 |
| BAMM_tv2 | 0.347 | <0.0001 | 2129.037 |
| BAMM_tv3 | 0.345 | <0.0001 | 2133.446 |
| DR | 0.364 | <0.0001 | 1804.268 |

**Time model (Richness~Col):**

| Colonization time | $R^2$ | P | Deviance |
|---|---|---|---|
| Basin mean | 0.326 | <0.0001 | 2283.367 |
| Basin median | 0.317 | <0.0001 | 2313.507 |

**Full model using mean colonization times (Richness~Div+Col):**

| Div rate | NetDiv+ ColTimes ($R^2$) | NetDiv+ ColTime (P) | NetDiv+ ColTime (Dev) | Dev.Time (%) | Dev.Div (%) | Time(%) ÷ Div(%) |
|---|---|---|---|---|---|---|
| BAMM_tc1 | 0.370 | <0.0001 | 2045.578 | 8.896 | 3.217 | 2.766 |
| BAMM_tc2 | 0.367 | <0.0001 | 2054.468 | 8.564 | 3.675 | 2.331 |
| BAMM_tc3 | 0.367 | <0.0001 | 2054.772 | 8.552 | 3.776 | 2.265 |
| BAMM_tv1 | 0.371 | <0.0001 | 2041.522 | 9.048 | 3.306 | 2.737 |
| BAMM_tv2 | 0.374 | <0.0001 | 2031.353 | 9.428 | 3.655 | 2.580 |
| BAMM_tv3 | 0.375 | <0.0001 | 2029.326 | 9.504 | 3.895 | 2.440 |
| DR | 0.402 | <0.0001 | 1677.830 | 22.654 | 4.730 | 4.789 |

**Full model using median colonization times (Richness~Div+Col):**

| Div rate | NetDiv+ ColTimes ($R^2$) | NetDiv+ ColTime (P) | NetDiv+ ColTime (dev) | Dev.Time (%) | Dev.Div (%) | Time(%) ÷ Div(%) |
|---|---|---|---|---|---|---|
| BAMM_tc1 | 0.366 | 8.20E-26 | 2056.643 | 9.610 | 2.803 | 3.429 |
| BAMM_tc2 | 0.524 | 1.90E-21 | 1508.815 | 30.105 | 5.512 | 5.461 |
| BAMM_tc3 | 0.364 | 1.52E-25 | 2062.363 | 9.396 | 3.379 | 2.781 |
| BAMM_tv1 | 0.523 | 7.58E-22 | 1512.857 | 29.954 | 5.567 | 5.381 |
| BAMM_tv2 | 0.364 | 1.67E-25 | 2062.905 | 9.376 | 3.472 | 2.701 |
| BAMM_tv3 | 0.525 | 1.49E-21 | 1508.096 | 30.132 | 5.544 | 5.435 |
| DR | 0.529 | 2.03E-20 | 1494.228 | 30.651 | 5.017 | 6.110 |

**Table S5.** Results of spatially explicit Generalized Additive Regression models fitted to test the contributions of diversification rates and colonization times in driving species richness differences among basins. Here we also controlled for the simultaneous effects of surface area on richness by adding area as a covariate in the model. For each model, we indicate the corresponding $R^2$, P-value, and deviance. For bivariate models, we indicate the deviance explained by each predictor (dev.x%) along with the ratio of this index between variables (e.g. Time(%) ÷Div(%)). The larger this ratio is, the greater information lost compared to the full model by excluding time relative to diversification rates. Results are presented for 7 estimates of diversification rates and two different values of colonization times.

**Null model (Richness ~ 1):**

| $R^2$ | P | Deviance |
|---|---|---|
| 0.257 | <0.0001 | 2672.931 |

**Div model (Richness~Div):**

| Div rate estimate | $R^2$ | P | Deviance |
|---|---|---|---|
| BAMM_tc1 | 0.5404 | <0.0001 | 1492.7693 |
| BAMM_tc2 | 0.5374 | <0.0001 | 1502.7984 |
| BAMM_tc3 | 0.5371 | <0.0001 | 1503.8292 |
| BAMM_tv1 | 0.5349 | <0.0001 | 1510.7786 |
| BAMM_tv2 | 0.5371 | <0.0001 | 1503.6893 |
| BAMM_tv3 | 0.5342 | <0.0001 | 1513.6581 |
| DR | 0.5544 | <0.0001 | 1261.0086 |

**Time model (Richness~Col):**

| Colonization time | $R^2$ | P | Deviance |
|---|---|---|---|
| Basin mean | 0.534 | <0.0001 | 1573.977 |
| Basin median | 0.527 | <0.0001 | 1597.288 |

**Full model using mean colonization times (Richness~Div+Col):**

| Div rate | NetDiv+ ColTimes ($R^2$) | NetDiv+ ColTime (P) | NetDiv+ ColTime (Dev) | Dev.Time (%) | Dev.Div (%) | Time(%) ÷ Div(%) |
|---|---|---|---|---|---|---|
| BAMM_tc1 | 0.557 | <0.0001 | 1433.111 | 2.39 | 1.725 | 2.393 |
| BAMM_tc2 | 0.555 | <0.0001 | 1441.068 | 2.15 | 1.787 | 2.152 |
| BAMM_tc3 | 0.554 | <0.0001 | 1442.016 | 2.12 | 1.789 | 2.123 |
| BAMM_tv1 | 0.552 | <0.0001 | 1449.048 | 1.9 | 1.786 | 1.904 |
| BAMM_tv2 | 0.554 | <0.0001 | 1443.277 | 2.08 | 1.747 | 2.080 |
| BAMM_tv3 | 0.553 | <0.0001 | 1445.533 | 2.01 | 1.981 | 2.011 |
| DR | 0.577 | <0.0001 | 1182.997 | 4.58 | 2.398 | 4.578 |

**Full model using median colonization times (Richness~Div+Col):**

| Div rate | NetDiv+ ColTimes ($R^2$) | NetDiv+ ColTime (P) | NetDiv+ ColTime (dev) | Dev.Time (%) | Dev.Div (%) | Time(%) ÷ Div(%) |
|---|---|---|---|---|---|---|
| BAMM_tc1 | 0.554 | <0.0001 | 1442.629 | 2.785 | 1.441 | 1.932 |
| BAMM_tc2 | 0.552 | <0.0001 | 1450.271 | 2.556 | 1.512 | 1.691 |
| BAMM_tc3 | 0.551 | <0.0001 | 1451.600 | 2.517 | 1.504 | 1.674 |
| BAMM_tv1 | 0.550 | <0.0001 | 1456.117 | 2.381 | 1.579 | 1.508 |
| BAMM_tv2 | 0.552 | <0.0001 | 1450.320 | 2.558 | 1.537 | 1.665 |
| BAMM_tv3 | 0.551 | <0.0001 | 1453.464 | 2.464 | 1.727 | 1.427 |
| DR | 0.568 | <0.0001 | 1209.045 | 4.341 | 1.490 | 2.913 |

**Table S6.** Results of spatially explicit Generalized Additive Models testing the relationship between colonization times and diversification rates. Here we show the $R^2$ value of the regression model and the associated P-value. We also present rho and P values for the analyzed non-parametric correlation analyses between colonization times and diversification rates.

| **Predictor** | **Mean colonization** | | | | **Median colonization** | | | |
|---|---|---|---|---|---|---|---|---|
| | $R^2$ | P | rho | P-rho | $R^2$ | P | rho | P-rho |
| BAMM_tc1 | 0.393 | <0.0001 | -0.125 | <0.0001 | 0.393 | <0.0001 | -0.125 | <0.0001 |
| BAMM_tc2 | 0.395 | <0.0001 | -0.129 | <0.0001 | 0.395 | <0.0001 | -0.129 | <0.0001 |
| BAMM_tc3 | 0.394 | <0.0001 | -0.129 | <0.0001 | 0.394 | <0.0001 | -0.129 | <0.0001 |
| BAMM_tv1 | 0.390 | <0.0001 | -0.100 | <0.0001 | 0.390 | <0.0001 | -0.100 | <0.0001 |
| BAMM_tv2 | 0.390 | <0.0001 | -0.102 | <0.0001 | 0.390 | <0.0001 | -0.102 | <0.0001 |
| BAMM_tv3 | 0.390 | <0.0001 | -0.110 | <0.0001 | 0.390 | <0.0001 | -0.110 | <0.0001 |
| DR | 0.421 | <0.0001 | -0.104 | <0.0001 | 0.421 | <0.0001 | -0.104 | <0.0001 |

**Table S7.** Results of spatially explicit Generalized Additive Models testing for the effects of surface area in explaining latitudinal patterns in colonization times and diversification rates. Here we report the $R^2$ for the full model including latitude and area as predictors of colonization times and colonization times. Next, we indicate the percent of deviance in the full model that is independently explained by area (Area (%Dev.exp)) or latitude (Latitude (%Dev.exp)).

| **Response variable** | **Full model ($R^2$) and P** | **Full (Dev)** | **Area (%Dev.exp)** | **Area ($R^2$)** | **Latitude (%Dev.exp)** | **Latitude ($R^2$)** | **Latitude(%) ÷ Area(%)** |
|---|---|---|---|---|---|---|---|
| Net diversification (12k_tv3) | 0.182 (P<0.001) | 4.957 | 3.324 | 0.074 | 49.196 | 0.125 | 14.800 |
| Mean colonization time | 0.510 (P<0.001) | 22.886 | 0.587 | 0.025 | 70.082 | 0.388 | 119.390 |

**Table S8:** Results of models testing the relationship between the number of colonizations and basin richness, latitude, mean diversification rate, or mean colonization time. We estimated the number of independent regional colonizations represented among each basin's fauna. A visual representation of the models is presented in Figure S4.

| **Variable** | **Linear regression** | | | **Spatially explicit GAM** | |
|---|---|---|---|---|---|
| | Slope | $R^2$ | P-value | $R^2$ | P-value |
| Richness | 1.127 | 0.622 | <0.001 | 0.665 | <0.001 |
| Diversification rate (12k_tc3) | 0.001 | <0.001 | 0.1487 | 0.005 | 0.010 |
| Colonization time (mean) | -9.995 | 0.067 | <0.001 | 0.089 | <0.001 |
| Latitude | -0.001 | <0.001 | 0.1121 | 0.015 | <0.001 |

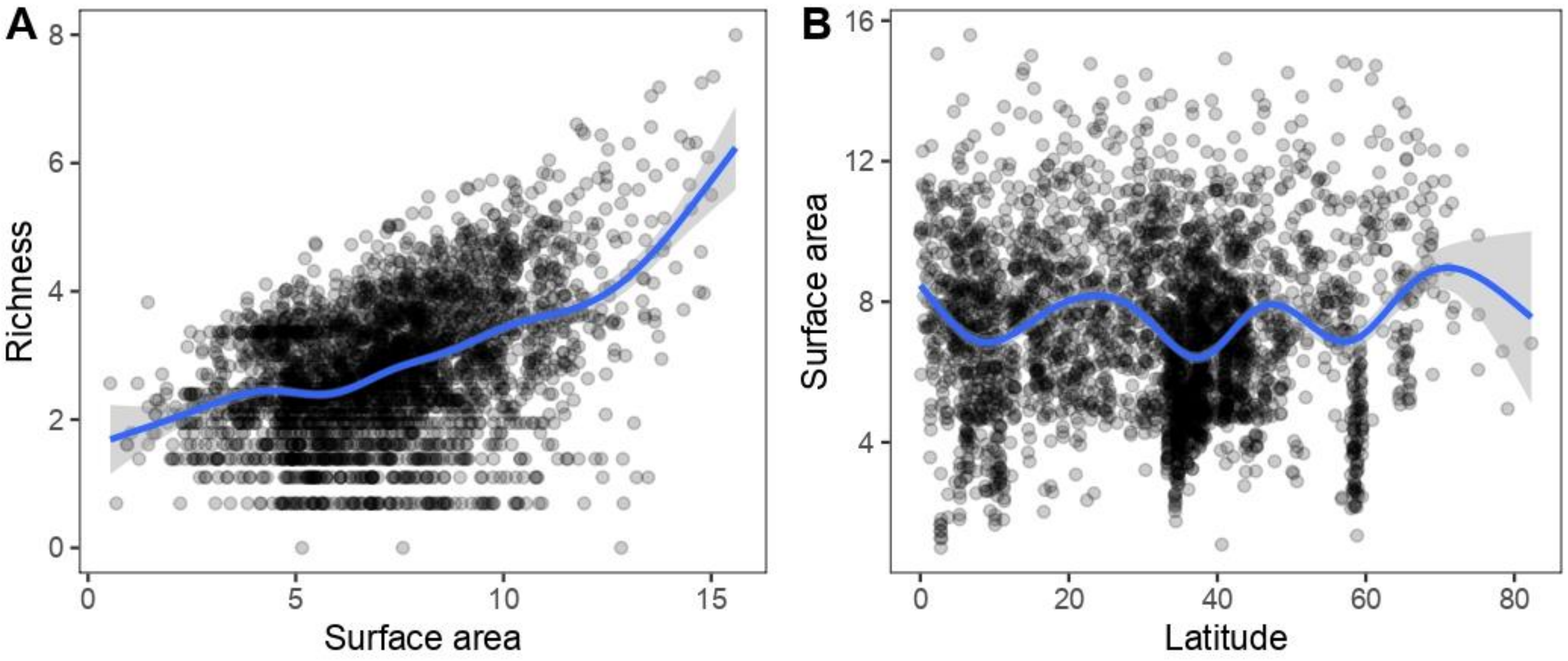


**Figure S1.** Scatterplots depicting the relationship between (A) surface area and richness, and (B) surface area and latitude. A GAM between richness and surface area was significant with $R^2$=0.242 and P<0.001. However, a GAM indicated that surface area and latitude are only weakly related ($R^2$=0.054, P<0.0001; Table S3).

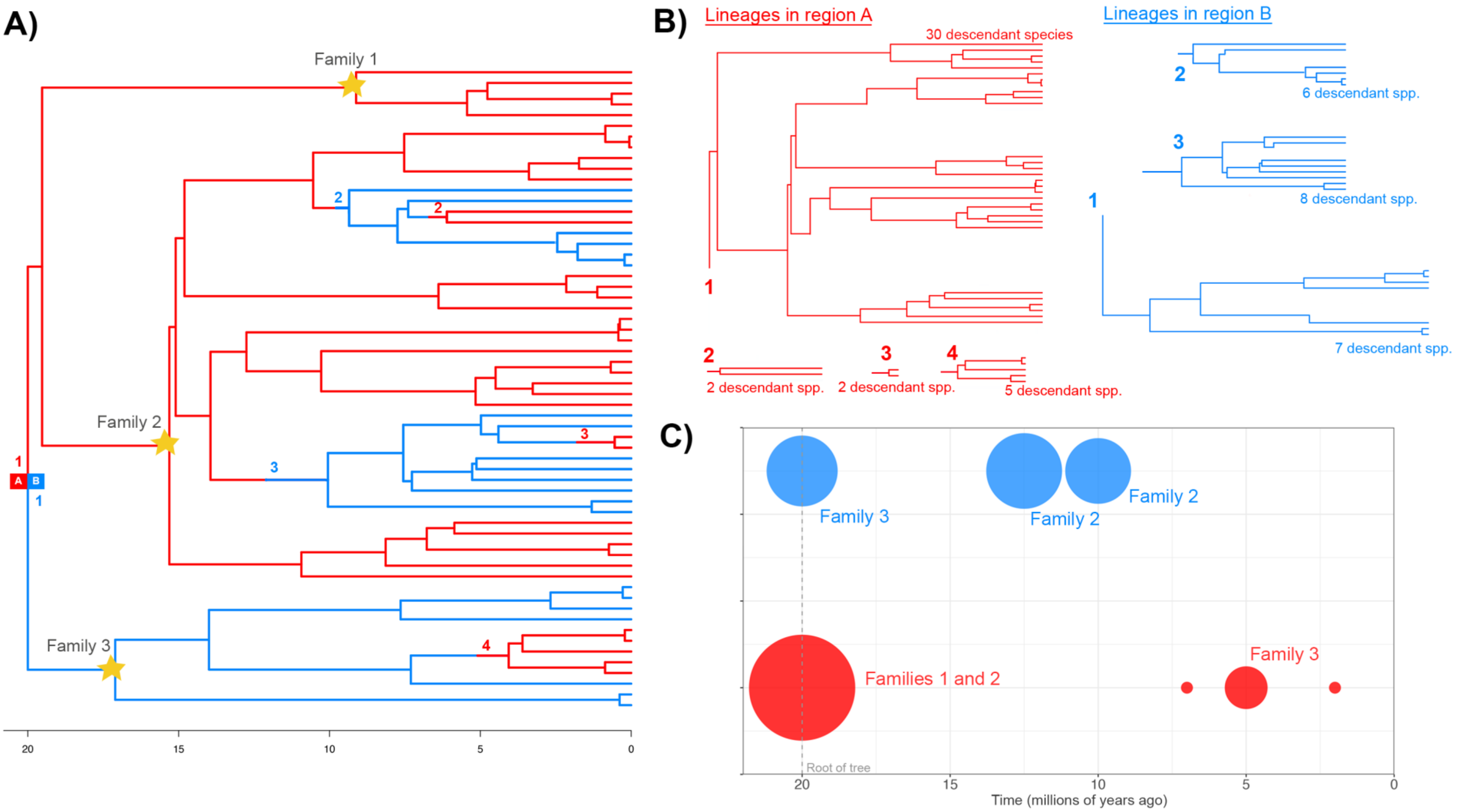
A)
Family 1
Family 2
Family 3
1
A
B
1
2
2
3
3
4
20
15
10
5
0
B)
Lineages in region A
30 descendant species
1
2
2 descendant spp.
3
2 descendant spp.
4
5 descendant spp.
Lineages in region B
2
6 descendant spp.
3
8 descendant spp.
1
7 descendant spp.
C)
Family 3
Family 2
Family 2
Families 1 and 2
Family 3
Root of tree
20
15
10
5
0
Time (millions of years ago)

**Figure S2.** An approach to visualizing complex relationships between colonization, time and richness (see Figure 5 in main text). This approach is designed for phylogenies that are too large to display effectively. (A) A hypothetical order is distributed across two regions A (red) and B (blue) and contains three families with crown ages marked by a star. Ancestral area reconstructions on a time-calibrated phylogeny are used to identify lineages that dispersed to new regions. Note that occupancy in a region often precludes the origin of named clades such as families. We used this approach to identify the colonization times used in GAM analyses. (B) For further visualization, lineages representing unique colonizations are tallied, along with the number of living descendants. Nested lineages are excluded from this tally if they left the focal region. As such, groups associated with unique colonizations are often paraphyletic. (C) A bubble plot displays lineages occupying each region by the timing of their arrival to that region. The bubbles are sized in scale of the proportion of living species in the region descended from each lineage. Dispersal to new regions can occur at any time along the phylogeny, and so these lineages often do not correspond to named clades. Still, recognizable clade names are added for context, referring to the living descendants of these colonization events.

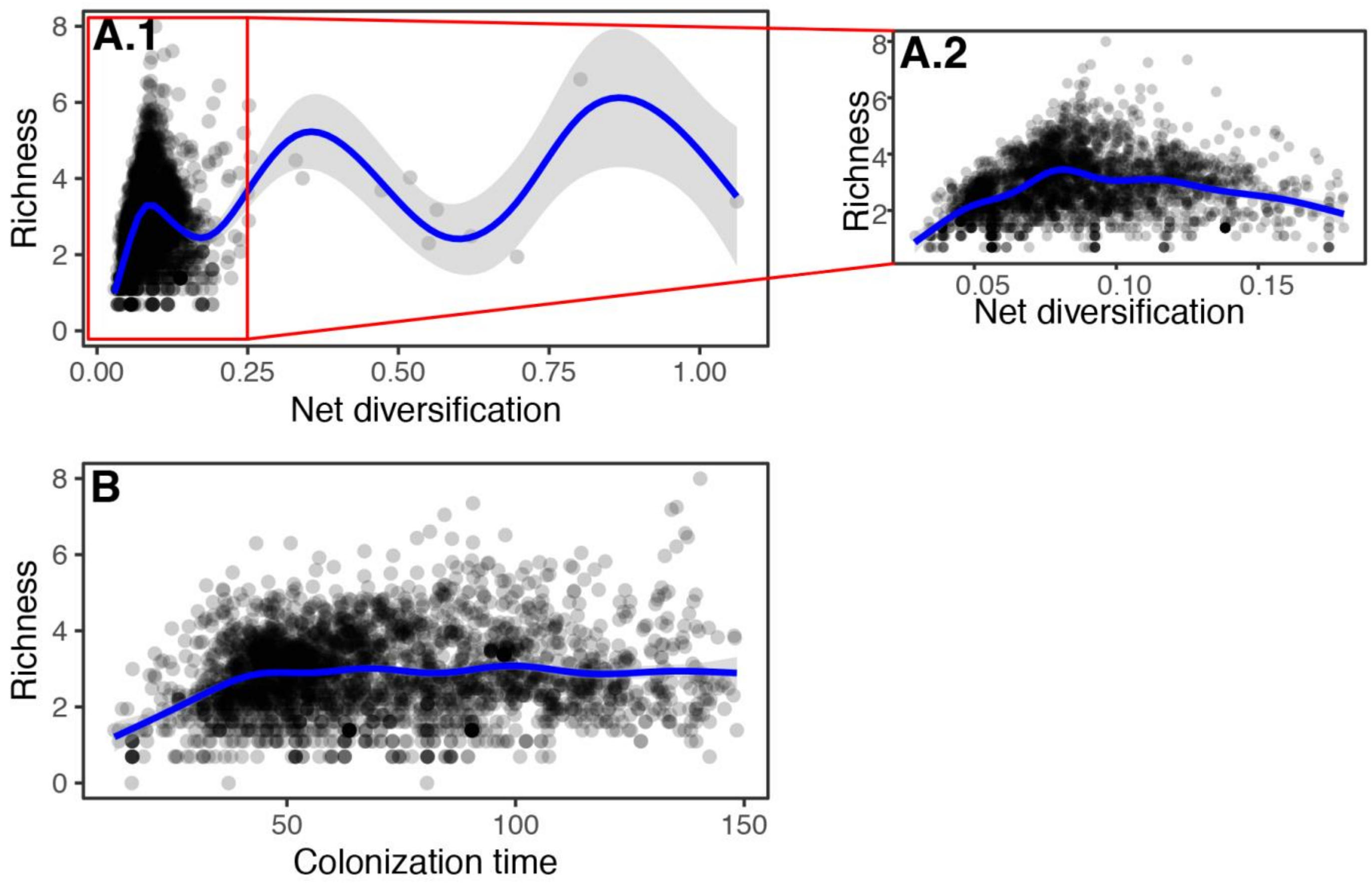


**Figure S3.** Scatterplot depicting the relationships between basin richness vs. net diversification (**A.1**; both log-transformed; GAM, $R^2$=0.346, P<0.001, deviance explained=3.217%; ; Table S4), and richness vs. mean colonization time (**B;** GAM, $R^2$=0.326, P <0.001, deviance explained=8.896%; Table S4). Inset (**A.2**) shows the trend after excluding 47 exceptionally fast-diversifying basins (basins with rates above 1.025 events/My including 27 in the Afrotropics and three in the Neotropics). Diversification rates shown here were estimated using BAMM under a time-constant rates model and represent the mean tip-associated values of species found in each basin (Rabosky et al., 2018). Colonization of major regions was inferred from ancestral area reconstructions (Matzke, 2014); points represent the mean time of colonization associated with species in each basin. See Table S4 for quantitative estimates of the regression models.

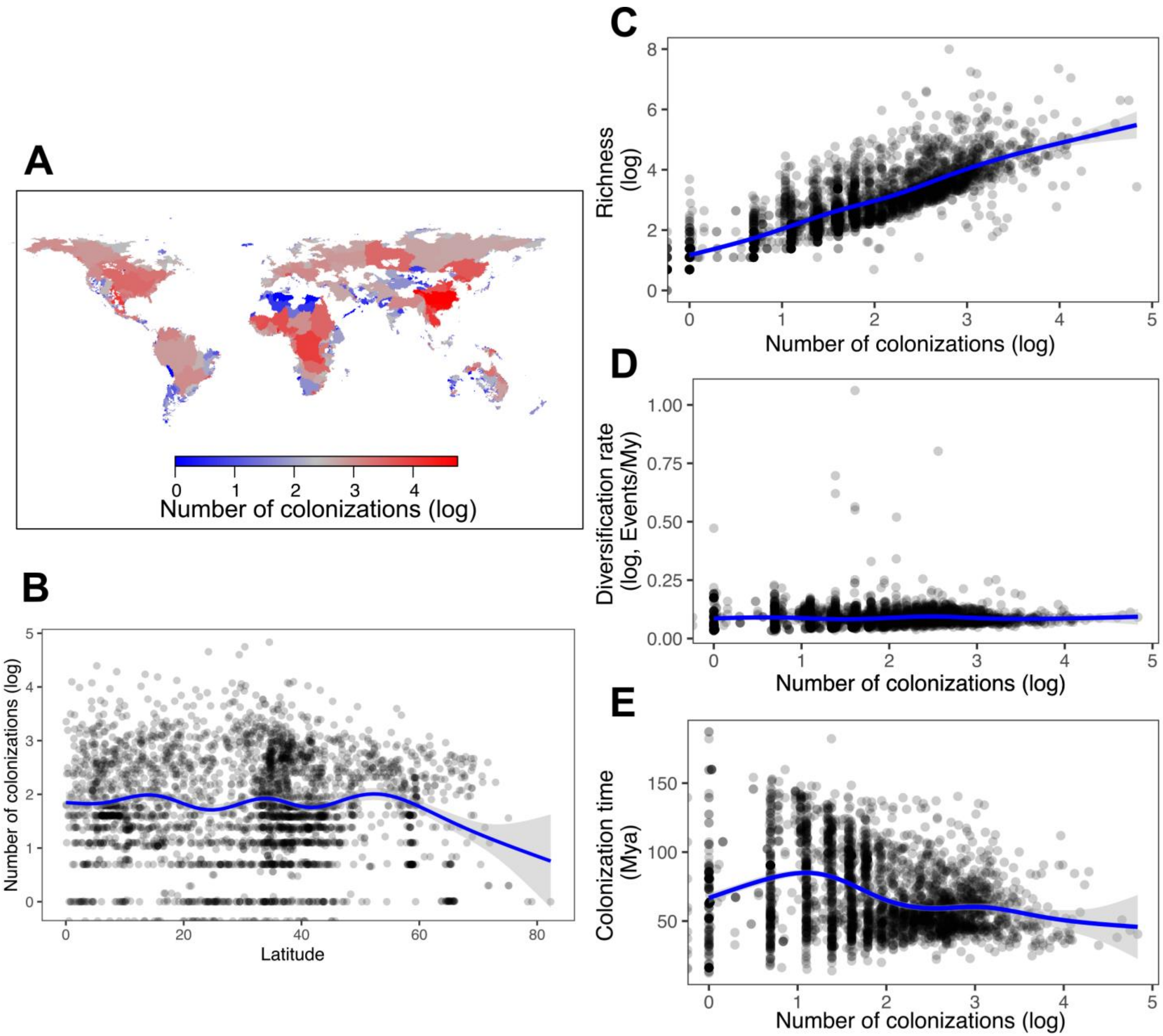

A
0
1
2
3
4
Number of colonizations (log)
B
Number of colonizations (log)
Latitude
0
20
40
60
80
C
Richness (log)
Number of colonizations (log)
D
Diversification rate (log, Events/My)
1.00
0.75
0.50
0.25
0.00
Number of colonizations (log)
E
Colonization time (Mya)
150
100
50
Number of colonizations (log)

**Figure S4.** (**A**) The spatial context of the number of regional colonizations. For each basin, we added the number of independent colonizations of the region represented among species in that basin. Values shown here are the mean counts among 100 biogeographic stochastic maps. (**B**–**E**) Scatterplots depicting the relationships between the number of colonizations and latitude (**B**; linear model, slope=-0.001, $R^2$=0.015, P=0.1121), basin richness (**C**; slope=1.127, $R^2$=0.622, P<0.001), diversification rates (**D;** slope=0.001, $R^2$<0.001, P=0.148), or mean colonization times (**E;** slope=-9.995, $R^2$=0.067, P<0.001). Diversification rates shown here were estimated using BAMM under a time-constant rates model and represent the mean tip-associated values of species found in each basin (Rabosky et al., 2018). Full results are presented in Table S8, along with results using spatially-explicit GAMs to describe the same set of relationships.

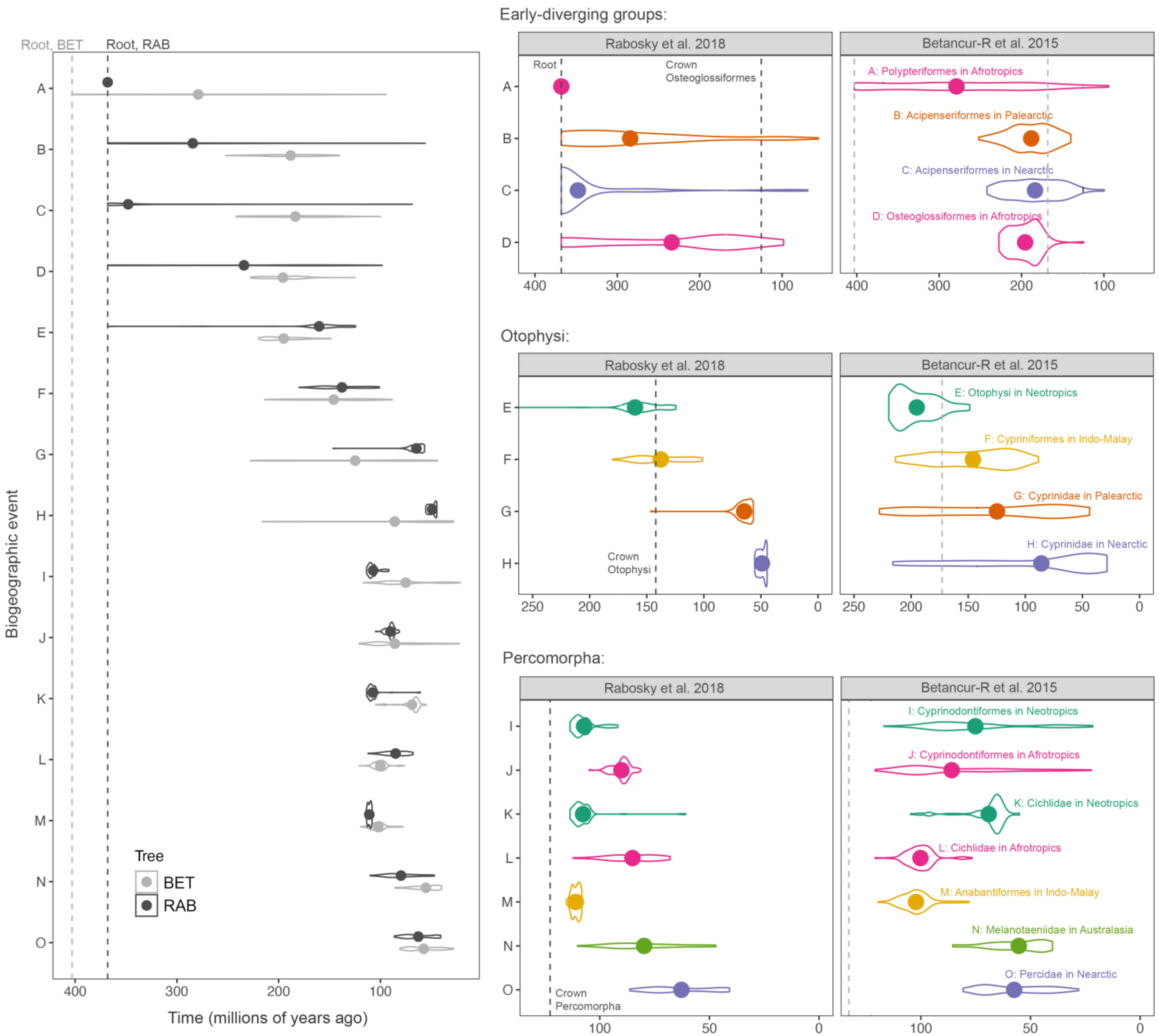
Root, BET
Root, RAB
Biogeographic event
Time (millions of years ago)
Tree
BET
RAB
Early-diverging groups:
Rabosky et al. 2018
Betancur-R et al. 2015
Root
Crown Osteoglossiformes
A: Polypteriformes in Afrotropics
B: Acipenseriformes in Palearctic
C: Acipenseriformes in Nearctic
D: Osteoglossiformes in Afrotropics
Otophysi:
Crown Otophysi
E: Otophysi in Neotropics
F: Cypriniformes in Indo-Malay
G: Cyprinidae in Palearctic
H: Cyprinidae in Nearctic
Percomorpha:
Crown Percomorpha
I: Cyprinodontiformes in Neotropics
J: Cyprinodontiformes in Afrotropics
K: Cichlidae in Neotropics
L: Cichlidae in Afrotropics
M: Anabantiformes in Indo-Malay
N: Melanotaeniidae in Australasia
O: Percidae in Nearctic

**Figure S5.** Comparing the timing of important biogeographic events inferred using two phylogenies of ray-finned fishes. The molecular phylogeny from Rabosky et al. (2018; "RAB") was used in our main analyses and contains 11,499 species (all extant). The phylogeny from Betancur-R et al. (2015; "BET") contains 240 fossils and 1,582 living species. In each case, we fit a DEC model and performed biogeographic stochastic mapping. Violin plots display the probability density of dates for key events (first arrivals of important clades to regions) inferred from 100 simulated histories. Solid points represent the mean dates among 100 simulations (these mean dates were used in our GAM analyses). Note that the occupancy of the region often precedes the crown age of the clade (see right panels and Figure S2). The left panel shows the correspondence between the two phylogenies for inferred dates. The right three panels show a close-up of the date range for groups of similar age (early-diverging, otophysan, and percomorph lineages) inferred using each tree. For detailed discussion see Extended Results 1.
